# Modulation Of SUDEP By Central Serotonergic Cooperating with Noradrenergic Circuits: A Synergistic-Dependent Manner

**DOI:** 10.1101/2024.07.20.604434

**Authors:** Qing Xu, XiaoXia Xu, YaXuan Wu, Yue Yang, LeYuan Gu, ZhuoYue Zhang, ZiWen Zhang, XuanYi Di, YuLing Wang, Qian Yu, XiTing Lian, HaiXiang Ma, HongHai Zhang

## Abstract

Sudden unexpected death in epilepsy (SUDEP) is the leading cause of death in refractory epilepsy patients. Despite previous accumulating evidence has shown that seizure-induced respiratory arrest (S-IRA) may play the main contributor to SUDEP as an initiating event preeminent cause of mortality, the specific underlying mechanism of action remains unclear. Based on our previous work, serotonin (5-HT) signaling in the dorsal raphe nucleus (DRN) is strongly implicated in S-IRA in animal models, including the DBA/1 mice, on the meanwhile, norepinephrine (NE) neurons of the locus coeruleus (LC) also plays a vital role in regulating respiratory function on its own. Superficially, monoaminergic neuron, as important neurotransmitters in the central nervous system, have similar modes of action in the maintenance of nervous system balance, and each of them has a regulatory effect on SUDEP. However, it remains to be investigated whether monoaminergic neuron family (NE and 5-HT) are related in the mechanism of regulating SUDEP, what is even more curious is whether the two are intrinsically linked. Thus, we hypothesize neural mechanism of central noradrenergic and serotonergic circuits in modulating SUDEP in a synergistic-dependent manner, this endeavor will culminate in a significant breakthrough in elucidating the precise mechanism of action underlying SUDEP. In our study, we will use chemogenetics, optogenetics, calcium signal recording, and bidirectional tracing to explore the internal mechanism of DR-LC regulating the occurrence of SUDEP, and by specifically injecting 5-HT2AR antagonist Ketanserin (KET) and/or NEα-1R antagonist Prazosin into the pre-Bötzinger complex (PBC), it was finally elucidate that the DR-LC-PBC network can effectively reduce the incidence of SIRA. We firstly proposed a powerful target for exploring the reduction of the incidence of SUDEP, which has great clinical translation potential.

## 1 Introduction

Epilepsy stands as the second most prevalent neurological disorder, trailing only behind stroke, with a global prevalence of approximately 70 million individuals^1,2^. Recent investigations have highlighted that the risk of SUDEP in pediatric patients is approaching that of adults, while the likelihood escalates during pregnancy among epileptics^3^. Furthermore, SUDEP emerges as the primary contributor to mortality among young adults with epilepsy, exhibiting an incidence rate two to three times higher than other conditions leading to sudden death in this demographic^4^. Notably, only approximately one-third of these patients receive adequate or sufficient therapeutic intervention. Sudden Unexpected Death in Epilepsy (SUDEP) represents a highly critical yet frequently overlooked complication among this patient population, constituting the foremost direct cause of premature mortality associated with epilepsy^5^. The precise mechanisms underlying SUDEP remain elusive, though they are postulated to involve intricate interactions among autonomic nervous system dysfunction, alterations in brainstem function, cardiorespiratory failure, and widespread suppression of postictal electroencephalographic activity. Among the identified risk factors, generalized tonic-chronic seizures (GTCS) and nocturnal seizures stand out as pivotal^6,7^. Specifically, the neuronal networks activated during GTCS may impair brainstem respiratory or autonomic control centers, triggering hypoventilation, apnea, and cardiovascular collapse, ultimately culminating in patient demise. Recent scholarly endeavors have increasingly emphasized the significance of apnea and cardiac dysfunction as potential pivotal mechanisms in SUDEP. In particular, seizure-induced respiratory arrest (S-IRA) has emerged as a primary precipitating factor in numerous fatal cases^7–9^. Consequently, there is an urgent need within the research community to elucidate the intricate mechanisms of SUDEP and devise strategies to mitigate or avert its occurrence, thereby offering tangible clinical translational benefits.

5-Hydroxytryptamine (Serotonin, 5-HT,) plays a pivotal role in the central nervous system (CNS), encompassing the regulation of appetite, sleep, memory, learning, emotion, and behavior^10,11^. Notably, 5-HT emerges as a potential key player in preventing the onset and progression of Sudden Unexpected Death in Epilepsy (SUDEP)^12^. Studies suggest that 5-HT reuptake inhibitors can mitigate S-IRA (seizure-induced respiratory arrest) in DBA/1 mice triggered by audiogenic seizures by elevating 5-HT levels in the synaptic cleft^13^. The Dorsal Raphe Nucleus (DRN), the primary source of forebrain 5-HT innervation, harbors 5-HT neurons whose activity is intimately linked to epilepsy^14,15^. In the preliminary stages of our research, the group has made a noteworthy observation in a DBA/1 mouse model of SUDEP: optogenetic stimulation of DRN 5-HT neurons has been found to markedly diminish the occurrence of S-IRA, indicating a promising therapeutic avenue^16^. Norepinephrine (NE), as another crucial neurotransmitter, is synthesized and secreted primarily by postganglionic sympathetic neurons and noradrenergic neurons within the brain, with the latter responsible for its release. The Locus Coeruleus (LC), a major NE-synthesizing and releasing nucleus in the brain, projects noradrenergic fibers to the Nucleus Tractus Solitarius (NTS), a vital center for respiratory and circulatory control, forming synaptic connections^17–19^. Our previous findings indicate that enhancing peripheral and central NE levels markedly decreases SUDEP incidence in DBA/1 mice, and with LC-NE neurons being of paramount importance, and deficient synthesis of NE and norepinephrinergic neurotransmission contributed to S-IRA and that the NEα-1R is a potential therapeutic target for the prevention of SUDEP^20,21^. Both 5-HT and NE, as major neurotransmitters within the brain and vital hormones in peripheral circulation, participate in regulating respiration, cardiac activity, sleep-wake cycles, and the autonomic nervous system, encompassing both sympathetic and parasympathetic components^22^. These neurotransmitters are intimately associated with S-IRA and SUDEP. However, as members of the catecholamine family, their specific mechanism of action is still unknown. We wanted to know whether 5-HT and NE are intrinsically connected when they play the same role, and whether there are common pathways and bridges of action. Based on the above studies, we discovered another nucleus known for its role in regulating respiratory rhythms, the pre-Bötzinger complex (PBC), and we observed that the seraeric neural circuit between DR and PBC was involved in the acoustic stimulation of DBA/1 mice or the S-IRA induced by pentadiazole (PTZ) injection, and found that 5-HT2AR, located in PBC, plays an important role in this^23^.

Given DRN and LC pivotal roles as central regulators of respiratory and circulatory functions and as key inhibitors of SUDEP, this study employs sound-induced and pentylenetetrazol (PTZ)-induced SUDEP models to explore the mechanisms and interactions between 5-HT and noradrenergic neurons in modulating SUDEP^24^. We initially administered 5-Hydroxytryptophan (5-HTP), the norepinephrine reuptake inhibitor Atomoxetine, and the Selective Serotonin-Norepinephrine Reuptake Inhibitor (SNRIs) Venlafaxine to DBA/1 mice via intraperitoneal injection, aiming to elevate 5-HT and NE levels and observe their effects on S-IRA^20^. To further delve into the specific receptor targets mediating these neural systems’ regulation of SUDEP, we injected the 5-HT2AR antagonist Ketamine (KET) and the NE α-1R antagonist Prazosin, discovering that they reversed the S-IRA-reducing effects of Venlafaxine. Employing in vivo neurophysiological techniques, including calcium signaling fiber photometry and optogenetic modulation, we monitored the activity changes of 5-HT neurons in the DRN and NE neurons in the LC. Furthermore, we specifically injecting KET and/or Prazosin into the pre-Bötzinger complex (PBC), discovering that they reversed the S-IRA-reducing effects of Venlafaxine. Our findings underscore the intricate interplay between these three systems, all of them playing crucial roles in regulating SUDEP. This study not only highlights the collaborative regulation of S-IRA and SUDEP by the DRN and LC systems but also suggests the clinical translational potential of PBC in treating epilepsy patients with S-IRA, thereby preventing SUDEP.

## 2 Materials and methods

### 2.1 Animals

All experimental procedures adhered to the Guidelines for the Care and Use of Laboratory Animals issued by the National Institutes of Health, which were approved by the Animal Advisory Committee of Zhejiang University. Proper animal use protocols and institutional guidelines were strictly followed. The DBA/1 mice used in this study were 7-8 weeks old, weighing 15-20g, and were purchased from Vital River Laboratories, SLAC Laboratory Animal Co., Ltd., and Jihui Laboratory Animal Co., Ltd. They were housed in a specific pathogen-free (SPF) barrier facility at the Animal Center of the School of Medicine, Zhejiang University, with ad libitum access to water and food, and maintained under optimal temperature and humidity conditions. Given the sensitivity of DBA/1 mice to sound, the housing environment was kept quiet to avoid excessive noise levels, preserving their auditory sensitivity. Prior studies have shown that gender does not affect the incidence rate in DBA/1 mice; hence, both male and female DBA/1 mice were selected for this study based on breeding availability^25^.

The TH-Cre transgenic mice employed in this experiment were generated through multiple rounds of crossing and backcrossing between TH-Cre transgenic mice on a C57BL/6J background and DBA/1 mice, resulting in purified DBA/1 background TH-Cre transgenic mice. The C57BL/6J background TH-Cre transgenic mice were sourced from Jackson Lab (strain B6.Cg-7630403G23RikTg(TH-Cre)1Tmd/J), with a genotype of Hemizygous for 7630403G23RikTg(TH-Cre)1Tmd serving as the Cre animal model. By performing marker-assisted accelerated backcrossing between hemizygous transgenic C57BL/6J TH-Cre mice and wild-type DBA/1 mice, hemizygous transgenic DBA/1 TH-Cre mice (hereinafter referred to as transgenic DBA/1 mice) were created. Congenic transgenic DBA/1 mice (with approximately 100% DBA/1 genetic background) were generated from the offspring of the fifth backcross generation. These hemizygous transgenic DBA/1 mice were then crossed with wild-type DBA/1 mice to produce the hemizygous transgenic DBA/1 mice used in the experiments.

### 2.2 PCR-based genotyping for TH-Cre transgenic mice

One week after birth, a 3-5mm segment of the toe or tail was clipped from the transgenic mice using alcohol-soaked scissors and placed into a sterile EP tube. The tube was then centrifuged to sediment the tissue at the bottom of the tube. A mixed solution of AD1/AD2 Buffer was prepared at a ratio of 4:1 (e.g., 10 µl of AD1 Buffer and 2.5µl of AD2 Buffer). This mixture was thoroughly combined and added to the centrifuge tube containing the tissue. After incubation at room temperature for 10 minutes, the mixture was heated at 95 °C for 3 minutes. Subsequently, AD3 Buffer was added, and the solution was thoroughly mixed. The mixture was then stored at 4 °C and used for PCR within one week. The PCR reaction mixture was set up as follows: 4µl of primers, 12.5µl of Taq enzyme, 7µl of ddH2O, and 1.5µl of the sample. The prepared samples were placed into a PCR instrument for reaction. Three grams of agarose were taken into a conical flask, and 100ml of TAE was added. The mixture was heated in a microwave oven for 2 minutes until boiling, then removed and thoroughly mixed. When the temperature dropped to 60°C, 10µl of dye was added, mixed again, and poured into a mold. Bubbles were removed, and an electrophoresis loading comb was inserted. After the gel had solidified, the electrophoresis comb was removed and samples were loaded. The electrophoresis apparatus was set to a constant voltage of 170V for 18 minutes. Following electrophoresis, the gel was developed, and bands were compared to screen for DBA/1 background TH-Cre mice for use in subsequent experiments.

### 2.3 Seizure induction and resuscitation

Auditory Stimulus-Induced Epilepsy Model: DBA/1 mice underwent continuous sound stimulation for 3-5 days between postnatal days 26-28 to establish audiogenic seizures and susceptibility to S-IRA (Sound-Induced Reflex Absence). The mice were placed in a cylindrical plexiglass container within a sound-proof chamber and subjected to acoustic stimulation using an electric bell (96dB SPL, Zhejiang People’s Electronics, China)) until the onset of audiogenic seizures (AGSz) and cessation of S-IRA (Sound-Induced Reflex Absence), with a maximum sound stimulation duration of 60 seconds. Following the occurrence of S-IRA (Sound-Induced Reflex Absence) in the mouse model of sound-stimulus induced SUDEP (Sudden Unexpected Death in Epilepsy), resuscitation was initiated within 5 seconds of the last respiratory gasp using a rodent ventilator. The ventilator parameters were set at 180 breaths per minute, with an inspiratory-to-expiratory (I/E) ratio of 1:1.5, and a tidal volume of 1cc of air. PTZ-Induced Epilepsy Model: DBA/1 mice aged 7-8 weeks, with a body weight of 15-20g, were selected and subjected to a single intraperitoneal injection of PTZ (Cat # P6500; Sigma-Aldrich, St. Louis, MO, USA) at 75 mg/kg to induce the occurrence of generalized seizures (GSz) and S-IRA^26^.

### 2.4 Pharmacological Intervention Experiments

All experiments were conducted in an animal laboratory with appropriate temperature, humidity, adequate lighting, and a quiet environment. Mice used in the experiments were mostly littermates, with an age difference between litters not exceeding one week. For 3-4 days prior to the start of the experiment, the mice cages were avoided from being changed, and the mice were gently stroked daily to allow them to become familiar with the experimenter’s scent and the environment, as well as to acclimate to being “picked up or held,” facilitating the subsequent connection of the optical fiber to their heads for the experiment. On the day of behavioral testing, the mice were placed in the behavioral testing room in advance to acclimate for 30-60 minutes, aiming to minimize nonspecific stress stimuli and the influence of other irrelevant factors on their activity levels and experimental results.

5-HTP (Cat #107751, Sigma-Aldrich, USA) is an intermediate metabolite of the essential amino acid L-tryptophan (LT) in the biosynthetic pathway of serotonin. 5-HTP exerts its effects primarily by enhancing the levels of serotonin within both the central and peripheral nervous systems. Clinically, it is often utilized as an antidepressant, appetite suppressant, and sleep aid^27^. Atomoxetine (Ca #Y0001586, Sigma-Aldrich, USA) is a selective norepinephrine reuptake inhibitor that targets and inhibits the presynaptic norepinephrine transporter (NET), thereby blocking the reuptake of NE into presynaptic vesicles and increasing the levels of NE in the synaptic cleft. It is currently used in the treatment of attention deficit hyperactivity disorder (ADHD) in children, adolescents, and adults^28^. Both 5-HTP and Atomoxetine were prepared using 0.9% saline solution.

#### 2.4.1 5-HTP and atomoxetine reverse effects on acoustic stimulation or PTZ-induced S-IRA in DBA/1 Mice

Twenty-four hours prior to the experiment, the susceptibility of DBA/1 mice that have undergone sound stimulation modeling to AGSz and S-IRA was confirmed.

In the acoustic stimulus model: the administration time for 5-HTP was set at 1 hour prior to the sound stimulation, where DBA/1 mice were administered with IP injections of either 5-HTP (100 mg/kg or 125 mg/kg, treatment group) or Saline (vehicle control group) (DB01-1-0701, Co., Ltd, China). For Atomoxetine, the administration time was 2 hours prior to the sound stimulation, with DBA/1 mice receiving IP injections of Atomoxetine (5, 10, 15, or 20 mg/kg, treatment group) or Saline (vehicle control group).

In the PTZ injection-induced model: DBA/1 mice were pre-treated with IP injections of either 5-HTP (100 mg/kg or 125 mg/kg) or Saline for 1 day. On the second day, 1 hour prior to the IP injection of PTZ (Cat #P6500, Sigma-Aldrich) at 75 mg/kg, the pre-treated mice were administered with IP injections of 5-HTP (100 mg/kg or 125 mg/kg) or Saline again. The administration time for Atomoxetine was set at 2 hours prior to the IP injection of PTZ (75 mg/kg), where DBA/1 mice were administered with IP injections of either Atomoxetine (5 mg/kg or 15 mg/kg) or Saline.

During both experiments, the behavioral activities of the mice throughout all the processes were video-recorded using cameras placed inside a sound-proof chamber. These videos were subsequently used for post-analysis to determine the Incidence of S-IRA, AGSz latency, Duration of W+C, Duration of tonic-clonic seizures, and Seizure score.

#### 2.4.2 Effect of venlafaxine on acoustic stimuli or PTZ model-induced S-IRA in DBA/1 Mice

Venlafaxine (PHR1736, Sigma-Aldrich, USA) is a serotonin (5-HT) and norepinephrine (NE) reuptake inhibitor, which blocks the reuptake of 5-HT and NE neurotransmitters. Clinically, Venlafaxine is primarily used for the treatment of major depressive disorder and generalized anxiety disorder. At low doses, Venlafaxine functions similarly to Selective Serotonin Reuptake Inhibitors (SSRIs), acting exclusively on 5-HT receptors. However, at medium to high doses, it acts on both 5-HT and NE receptors, while exhibiting low affinity for histamine receptors and cholinergic receptors^29^. Venlafaxine was prepared using 0.9% saline solution.

Twenty-four hours prior to the experiment, the susceptibility of DBA/1 mice that have undergone sound stimulation modeling to AGSz and S-IRA was confirmed. In the acoustic stimulus model: thirty minutes prior to the sound stimulation, DBA/1 mice were administered with IP injections of various doses of Venlafaxine (5, 15, 25, 50, 75, or 100 mg/kg, treatment group) or Saline(vehicle control group).

In the PTZ injection-induced model: thirty minutes prior to the IP injection of PTZ (75 mg/kg), DBA/1 mice were administered with IP injections of various doses of Venlafaxine (5, 15, 25, 50, 75, or 100 mg/kg) or Saline.

During both experiments, the behavioral activities of the mice throughout all the processes were video-recorded using cameras placed inside a sound-proof chamber. These videos were subsequently used for post-analysis to determine the Incidence of S-IRA, AGSz latency, Duration of W+C, Duration of tonic-clonic seizures, and Seizure score.

#### 2.4.3 Effect of PCPA/DSP-4 on acoustic stimulation or PTZ-induced venlafaxine-mediated S-IRA inhibition in DBA/1 Mice

KET (Cat #8006, Sigma-Aldrich, USA) is a selective serotonin receptor antagonist that acts on serotonin receptors in the body, influencing the function of the 5-HT system by inhibiting the activity of serotonin receptors. KET primarily acts on the 5-HT2A receptor, functioning as a 5-HT2A receptor antagonist. Prazosin (Cat #P7791,Sigma-Aldrich,USA), on the other hand, is an NE-α 1R antagonist that blocks the effect of norepinephrine on α 1-receptors, and is commonly used for the management and treatment of hypertension, benign prostatic hyperplasia, and post-traumatic stress disorder-related conditions^30^. Both KET and Prazosin were prepared using 25% Dimethylsulfoxide (DMSO).

Twenty-four hours prior to the experiment, the susceptibility of DBA/1 mice that have undergone acoustic stimulation modeling to Acoustic Geniculate Stimulation-induced seizures (AGSz) and Spontaneous Irregular Respiratory Activity (S-IRA) was confirmed. In the acoustic stimulus model: thirty minutes prior to the sound stimulation, DBA/1 mice were administered with an intraperitoneal (IP) injection of Venlafaxine at a dose of 25 mg/kg or Saline as a control. Fifteen minutes before the sound stimulation, the experimental group received an IP injection of KET at a dose of 20 mg/kg, while thirty minutes before the sound stimulation, they received an IP injection of Prazosin at a dose of 0.01 mg/kg. The control group, on the other hand, received IP injections of 25% Dimethylsulfoxide (DMSO) (SHBK2703, Sigma-Aldrich, USA) at both 15 minutes and 30 minutes prior to the sound stimulation.

In the PTZ injection-induced model: thirty minutes prior to the intraperitoneal (IP) injection of PTZ (Pentylenetetrazole) at a dose of 75 mg/kg, DBA/1 mice were administered with an IP injection of Venlafaxine at a dose of 25 mg/kg or Saline as a control. Similarly, the experimental mice received an IP injection of KET at a dose of 20 mg/kg fifteen minutes before the PTZ injection, and an IP injection of Prazosin at a dose of 0.01 mg/kg thirty minutes before the PTZ injection. The control group received IP injections of 25% Dimethylsulfoxide (DMSO) at both 15 minutes and 30 minutes prior to the PTZ injection.

During both experiments, the behavioral activities of the mice throughout all the processes were video-recorded using cameras placed inside a sound-proof chamber. These videos were subsequently used for post-analysis to determine the Incidence of S-IRA, AGSz latency, Duration of W+C, Duration of tonic-clonic seizures, and Seizure score.

#### 2.4.4 Effect of PCPA/DSP-4 on PTZ-induced venlafaxine-mediated S-IRA inhibition in DBA/1 Mice

Para-chlorophenylalanine (PCPA)(C3635, Sigma-Aldrich, USA) is a selective and irreversible inhibitor of Tryptophan Hydroxylase (TPH), the rate-limiting enzyme in the synthesis of 5-HT. PCPA can inhibit the synthesis of 5-HT by inhibiting tryptophan hydroxylation, significantly reducing the peripheral and central concentrations of 5-HT, and is commonly used in animal behavioral studies such as sleep-wake, analgesia, and epilepsy^31,32^. N-(2-chloroethyl)-N-ethyl-2-bromobenzylamine hydrochloride (DSP-4) (C8417, Sigma-Aldrich, USA) is a selective neurotoxin targeting the noradrenergic system in the locus coeruleus of the brain. It readily crosses the blood-brain barrier and undergoes cyclization into a reactive aziridine derivative, which accumulates in noradrenergic nerve terminals via the norepinephrine transporter. Within the nerve terminals, the aziridine derivative reacts with cellular components, resulting in the destruction of the nerve terminals^20,33^. Our previous studies have also found that IP injection of DSP-4 significantly reduces the number of noradrenergic neurons in the locus coeruleus (LC)^17^. Both PCPA and DSP-4 were prepared using 0.9% saline as the solvent.

PCPA Group: DBA/1 mice were pretreated with IP injections of either PCPA (800 mg/kg) or Saline, administered either as a single dose 1 day prior or daily for 5 consecutive days. On the final day, mice were given an additional IP injection of either PCPA (800 mg/kg) or Saline 2.5 hours before PTZ injection. Thirty minutes before PTZ injection, mice were also administered IP injections of Venlafaxine (25 mg/kg) or Saline. DSP-4 Group: DBA/1 mice were pretreated with a single IP injection of either DSP-4 (50 mg/kg) or Saline. Thirty minutes before IP injection of PTZ (75 mg/kg), the pretreated mice received an IP injection of either Venlafaxine (25 mg/kg) or Saline. The behavioral activities of the mice throughout the drug-induced epileptic process were video-recorded using cameras. These videos were subsequently used for post-analysis to determine the Incidence of S-IRA, AGSz latency, Duration of W+C, Duration of tonic-clonic seizures, and Seizure score.

### 2.5 Immunohistochemistry Experiment

After the experiment, the mice were anesthetized with an IP injection of 1% pentobarbital sodium (IP, 50 mg/kg). The chest was opened along the sternum and ribs to fully expose the heart. A perfusion cannula was inserted into the left ventricle of the mouse and secured in place. A small incision was made in the right atrium, and a syringe filled with Phosphate Buffered Saline (PBS) was used to perfuse the entire body through the left ventricle until the liver of the mouse turned white. Subsequently, perfusion was continued with 4% Paraformaldehyde (PFA) until the tail of the mouse became stiff and elevated, indicating rigor mortis. During perfusion, care was taken to avoid excessive speed to prevent the fluid from entering the pulmonary circulation. After perfusion, the mice were decapitated, and the brains were removed and immersed in 4% PFA for fixation. The brains were then placed in a 4°C refrigerator overnight (approximately 12-16 hours). Following fixation, the brains were transferred to 5 ml centrifuge tubes and immersed in 5 ml of 30% sucrose solution for dehydration for 24 hours, until the brains sank to the bottom of the tubes, indicating sufficient dehydration. Using a cryostat, the dehydrated brain tissue was sliced into 35 or 40 μm thick sections. The brain slices were placed in well plates and washed with PBS by placing them on a shaker (40-50 r/min) for 5 minutes, repeating this process three times. A blocking solution was prepared on ice, and 200 μl of blocking solution was added to each well. The plates were placed on a shaker (10 r/min) and incubated at room temperature for 2.5 hours. After incubation, the blocking solution was aspirated, and the primary antibody dilution was added. The plates were then placed on a shaker in a 4°C refrigerator and incubated overnight. After incubation with the primary antibody, the brain slices were washed with PBS on a shaker (40-50 r/min) for 10 minutes, repeating this process three times. The brain slices were then transferred to a secondary antibody dilution and incubated in the dark for 1 hour. After incubation with the secondary antibody, the brain slices were washed with PBS on a shaker (40-50 r/min) for 15 minutes, repeating this process three times. Finally, the brain slices were stained with DAPI for 7-10 minutes and mounted with 60% glycerol. The VS120 Virtual Slide System and Nikon A1R Confocal Microscope were used to capture fluorescent images. The images were then analyzed using ImageJ image processing software and NIS-Elements Viewer to count and analyze the number of immunofluorescently stained positive cells and their co-localization.

### 2.6 Stereotaxic and Viral Injection Experiment

#### 2.6.1 Stereotaxic localization of mouse brain

Before the experiment, surgical instruments were sterilized using an autoclave, the surgical area was cleaned with 75% ethanol, a microinfusion pump was connected, and a stereotaxic apparatus and dental drill were assembled. The mice were anesthetized with an IP injection of 1% pentobarbital sodium (IP, 50 mg/kg), and a heating pad was turned on throughout the surgical procedure to maintain normal body temperature. The hair on the mouse’s head was shaved, and the mouse was secured in a stereotaxic apparatus using ear bars and a nose clamp, ensuring that the respiratory tract remained unobstructed. To prevent irreversible damage to the mouse’s vision from bright light, erythromycin eye ointment was applied to the surface of the mouse’s eyeballs, followed by a covering of sterile cotton balls for protection. After disinfecting with alcohol, the skin on the mouse’s head was cut open, and the connective tissue on the surface of the skull was wiped with an alcohol-soaked cotton ball. Hydrogen peroxide-soaked cotton balls were then used to gently wipe the cranial sutures, better exposing the anterior and posterior fontanelles of the mouse skull. The mouse skull was leveled using the tip of a syringe needle, ensuring that the height difference between the anterior and posterior fontanelles and the left and right sides (±2.30 mm) did not exceed 0.03 mm. Using the 4th edition of the mouse brain atlas, “Paxinos and Franklin’s The Mouse Brain in Stereotaxic Coordinates,” the coordinates for the DRN and LC were determined. A cranial drill was used to make a hole approximately 0.5-1 mm in diameter above the target nucleus. During this process, care was taken to avoid tissue bleeding, as excessive bleeding could lead to intracranial hematoma compression or even death of the mouse.

#### 2.6.2 Intranuclear viral injection

Before loading the virus, the microsyringe was filled with liquid paraffin, and the connection between the syringe and the glass electrode was sealed with hot melt glue. The airtightness of the syringe was tested by observing whether any bubbles formed inside the syringe. The virus was aspirated and positioned above the drilled hole targeting the desired brain region. The needle was slowly lowered while observing the mouse’s condition and ensuring that the tip of the needle did not deviate from its intended direction. After reaching the target brain region, the needle was left in place for 10 minutes to allow the brain tissue to fully recoil. The injection speed was set to 40-50nl/min, and the injection volume was 100nl. During the microinjection, the microscope was used to closely observe whether the stratification line between the virus and liquid paraffin in the glass electrode was descending. After the injection, the needle was left in place for an additional 10 minutes before being slowly and evenly withdrawn to prevent the virus from spilling upwards as the needle was removed. After successful injection, the skull surface was wiped clean of blood and bone debris using a wet cotton ball soaked in saline to keep the area clean and moist. The mouse was then removed from the stereotaxic apparatus, and the scalp was sutured using absorbable surgical suture. A layer of erythromycin ointment was applied to prevent infection at the surgical site. The mouse was placed on a 37°C constant temperature electric heating blanket to wait for anesthesia recovery. Once the mouse regained spontaneous activity, it was returned to its cage. Anterograde tracing virus HSV-1 strain H129 (Herpes simplex virus-1 H129, HSV-1 H129) possesses the characteristic of anterograde trans-synaptic transport across multiple synapses, enabling it to label entire neural networks. It is capable of labeling not only connections between different brain regions within the central nervous system but also connections between peripheral and central nervous system regions. Following viral infection, it can cross one synapse within 24-36 hours, two synapses within 36-48 hours, and up to three synapses within 60-72 hours. It is important to note that HSV-1 H129 virus injection is toxic to mice, with a typical survival time of 3-5 days after injection. In this experiment, the HSV-1 H129 infection time was set at 36-48 hours. In contrast, the retrograde tracer virus CTB-555 can be taken up from axonal terminals and retrogradely transported to label neuronal cell bodies, with an infection time of 1-2 weeks. Both optogenetic and calcium-indicating viruses have infection times of 3-4 weeks.

#### 2.6.3 Optogenetic/calcium signaling fiber optic recording manipulation

Drilling holes were made to the left and right anterior, as well as left and right inferior, of the bregma. Screws were then inserted into these holes, with one-third of the screw protruding from the skull surface to ensure a secure connection. This setup facilitated the subsequent attachment of dental cement for stabilization. The cannula/fiber optic holder was used to secure the cannula/fiber, which was then moved to the position above the drilled hole according to the coordinates of the target nucleus. The cannula/fiber was slowly lowered, and attention was paid to the mouse’s condition and any potential deviations during the descent. To avoid damaging blood vessels overlying the DRN, the fiber optic was implanted with a 10° rightward tilt at the DRN location, reducing postoperative mortality rates. Upon reaching the target position, the skull surface was wiped clean of bone debris and blood using a cotton ball soaked in saline. This step is crucial to ensure that the cannula is securely fixed and less prone to detachment. A mixture of dental impression material and powder was used to fix the cannula/fiber and screws. When securing the cannula, it is essential that the screws are completely covered by the dental cement, leaving only the upper half of the cannula/fiber visible. After the dental cement has set, the cannula/fiber holder was carefully and slowly released. A cannula core and cap were inserted into the cannula to prevent tissue from blocking it in the future. Following the surgery, the mouse was placed on a heated blanket for recovery and returned to its cage once spontaneous activity was confirmed. The optical fiber was implanted two weeks after viral infection, and the optogenetic experiments were conducted one week after fiber implantation. For cannula implantation, pharmacological experiments were performed one week later by connecting the injection cannula and tubing to a microinjector.

The target brain region coordinates involved in this study are as follows: DRN: AP −4.55 mm, ML ± 0.44 mm, DV −2.75 mm, with a 10° right tilt; LC: AP −5.45 mm, ML ± 0.9 mm, DV −3.65 mm; Lateral Ventricle: AP −0.45 mm, ML −1.0 mm, DV −2.50 mm. The AAV viruses utilized in the experiments were as follows: For optogenetics: pAAV-TPH2 PRO-ChETA-EYFP-WPRES-PAS and pAAV-CAG-DIO-CHETA-EGFP, both produced by Shanghai Shengbo Biomedical Technology Co., Ltd. For calcium signaling: AAV2/9-mCaMKIIa-GCaMP6f-WPRE-pA and rAAV-DBH-GCaMP6m-WPRE-hGH pA, both manufactured by Wuhan Shumi Brain Science and Technology Co., Ltd. Upon completion of the experiments, the locations of viral expression and cannula/fiber implantation were verified. This involved checking if the viral fluorescent expression and cannula tips were indeed within the target brain regions. Following this confirmation, behavioral data analysis was conducted accordingly.

### 2.7 Optogenetic manipulation and fiber optic recording of calcium signaling

Two weeks after the injection of optogenetic/calcium-signaling viruses, the optical fiber ferrule pin was implanted. One week following fiber implantation, the laser was connected to the ferrule pin on the mouse’s head via an optical fiber patch cord to conduct optogenetic/calcium-signaling fiber recording experiments.

The specific principle of optogenetics involves the exogenous expression of light-sensitive proteins to control or detect neural activity. The genetic information of the light-sensitive protein is incorporated into an AAV vector and delivered to specific types of cells within the nervous system, resulting in the expression of specialized ion channels. Upon absorbing excitation light of specific wavelengths, light-sensitive proteins can open ion channels, selectively allowing the passage of cations or anions, which subsequently alters the membrane potential. This change in membrane potential activates or inhibits neural activity. In this experiment, a 465 nm diode laser was used for the optical fiber laser system. For the DR light stimulation group, the light parameters were set to 20-ms pulse duration, 20 Hz, and 15mW, with a total stimulation duration of 20 minutes. For the LC light stimulation group, DBA/1 background TH-Cre transgenic mice were utilized, and both LC sides were stimulated with light parameters of 20-ms pulse duration, 20 Hz, and 15mW, for a total stimulation duration of 25 minutes. In both groups, pentylenetetrazol (PTZ) was administered intraperitoneally at 15 minutes and 20 minutes, respectively, during the light stimulation period. Behavioral observations and recordings were conducted, and immunohistochemical analysis was performed to quantify neuronal activation within the two nuclei.

The specific principle of calcium fiber recording involves calcium imaging technology, which is a subset of optogenetics. This technique enables the detection of changes in Ca2+ concentration within cells or tissues. The variation in calcium concentration is converted into a fluorescent signal, thereby transforming cellular electrical activity into a recordable optical signal. Utilizing viral vectors as tools, genetically encoded Ca2+ indicators are introduced into a specific type of neuron, enabling the expression of these Ca2+ sensors within targeted brain regions. These sensors are designed to detect changes in Ca2+ concentration. Excitation light travels through the optical fiber patch cord and the ferrule pin implanted in the animal’s head, reaching the targeted brain region. This light excites the genetically encoded Ca2+ indicators, resulting in the emission of a green fluorescent signal. The intensity of the excited fluorescent signal can be collected by the optical fiber tip implanted within the animal’s brain. The collected fluorescence is then converted into an electrical signal by a sensor and transmitted to a recording system, allowing for real-time observation of calcium fluorescence activity in a population of neurons within the studied brain region. In this study, various concentrations of Venlafaxine (1.25 mg/ml, 2.5 mg/ml, 6.25 mg/ml, 12.5 mg/ml, and 25 mg/ml, 2.0 μl) or Saline (2.0 μl) were injected intracerebroventricularly (ICV) through a cannula, while simultaneously recording calcium signals. The time points were aligned with the onset of seizures, and the fluorescence intensity changes in 5-HT neurons within the DRN and NE neurons within the LC were expressed as ΔF/F0 = (F−F0)/F0, where F represents the current fluorescence intensity and F0 represents the baseline fluorescence intensity. The analysis results were presented as a heat map.

### 2.8 Chemogenetics

Prior to the initiation of experiments by three weeks, a 100nL volume of a 1:3 mixture of mTH-Cre-AAV/DBH-Cre and pAAV-EF1a-DIO-hM3D-mCherry viruses (viral titer: 5.00×10¹² vg/mL, sourced from Brain VTA Technology Co., Ltd., Wuhan, China) was microinjected into either the locus coeruleus (LC) or the ventrolateral preoptic area (VLPO). Subsequent to the completion of chemogenetic manipulations, immunohistochemical analysis was employed to ascertain TH-hM3Dq expression levels within the LC and VLPO. For neuronal activation utilizing the chemogenetic approach, Clozapine-N-oxide (CNO; HY-17366, MedChemExpress) was dissolved in saline and administered to mice via intraperitoneal injection, 20 minutes subsequent to midazolam injection. In this investigation, two concentrations of CNO (0.1 mg/kg and 0.2 mg/kg) were evaluated, ultimately determining 0.2 mg/kg as the optimal dose for subsequent intraperitoneal administrations.

### 2.9 Quantification and statistical analysis

Experimental data were presented as mean ± standard error of the mean (Mean ± SEM). Prior to data analysis, all experimental data underwent the Shapiro-Wilk test for normality and the Levene’s test for homogeneity of variance. These tests ensure that the statistical assumptions are met for subsequent analyses. Comparison between two groups: If the data conform to a normal distribution, the student’s t-test is applied, including the independent samples t-test (which assumes both variance homogeneity and normal distribution). In cases where the data do not meet the criteria for normal distribution, the Mann-Whitney U test or the Wilcoxon signed-rank test is utilized. For comparisons involving three or more groups: If the data conform to a normal distribution and the variances are homogeneous, a One-way ANOVA analysis is performed. A P-value of < 0.05 is considered statistically significant. GraphPad Prism TM 8.0 and SPSS version 23.0 were utilized for statistical analysis of the data.

## 3 Results

### 3.1 Co-enhancement of 5-HT and NE neurotransmission in the brain significantly reduced the incidence of S-IRA evoked by acoustic stimulation or PTZ injection, exhibiting a synergistic effect

In order to explore the role of 5-HT and NE neurotransmission in the occurrence and development of SUDEP, particularly building upon previous work to further elucidate their interactive effects, we injected intraperitoneally varying doses of 5-HTP and Atomoxetine respectively in the DBA/1 mice induced by acoustic stimulation or PTZ injection (Fig.1A-B, 1E-F). Our findings revealed that, a notable dose-dependent decrease in the incidence of S-IRA was observed when monotherapy with a high dose of 125mg/kg 5-HTP or 15-20mg/kg Atomoxetine in both acoustic stimulation and PTZ injection induced DBA/1 mice. In contrast, the application of lower doses of 5-HTP or Atomoxetine does not comprehensively exhibit the effects of suppressing SUDEP. Intriguingly, the combined medication of ineffective doses of 100mg/kg 5-HTP and 5mg/kg Atomoxetine significantly reduces the incidence of S-IRA, while also prolonging the latency of AGSz seizures, decreasing the duration of wild running and clonic seizures, the duration of tonic-clonic seizures and seizure scores, surpassing the effects of any single drug dose we applied (Fig.1C-D, 1G-H). This underscores the potential synergistic effect of concurrently enhancing 5-HT and NE neurotransmission, however, which specific structures in brain are involved in regulating SUDEP and the exact targets of action remain to be elucidated.

**Figure 1.**
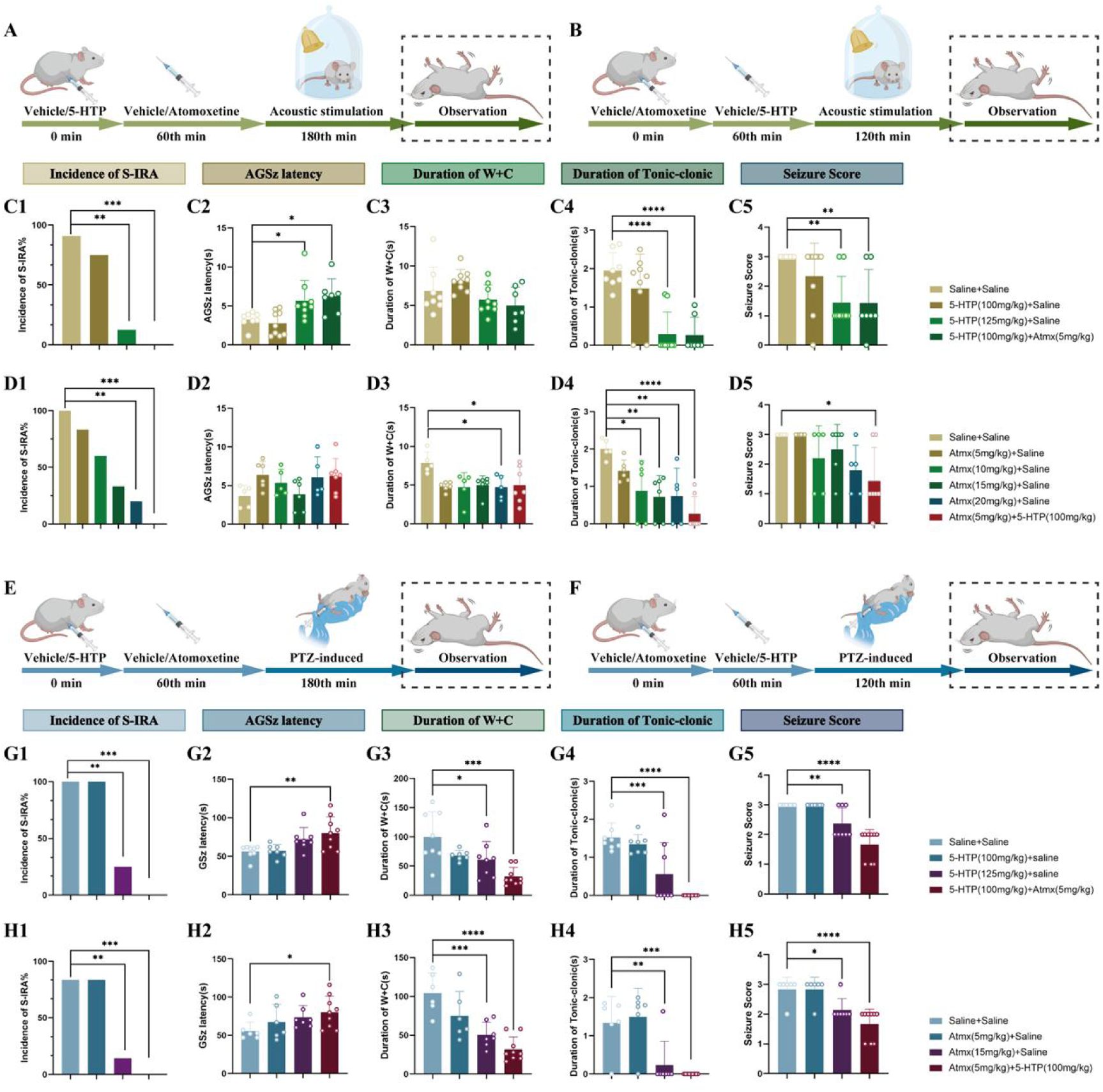
Elevated levels of 5-HT and NE demonstrate synergistic effects in reducing the incidence of S-IRA induced by acoustic stimulation and PTZ injection. **(A)** Schematic illustration of the observations in the DBA/1 mice induced by acoustic stimulation following IP injection of 5-HTP with or without Atomoxetine. **(B)** Schematic illustration of the observations in the DBA/1 mice induced by acoustic stimulation following IP injection of Atomoxetine with or without 5-HTP. **(C1-5)** The incidence of S-IRA evoked by acoustic stimulation, the duration of tonic-clonic seizures and the seizure score in the group with 125mg/kg 5-HTP and the group with 100mg/kg 5-HTP + 5mg/kg Atomoxetine were significantly reduced compared to the group with Saline. The AGSz latency in the group with 125mg/kg 5-HTP and the group with 100mg/kg 5-HTP + 5mg/kg Atomoxetine was markedly higher than the group with Saline. No obvious differences were observed between treatment groups in the duration of wild running and clonic seizures (*p* > 0.05). **(D1-5)** The incidence of S-IRA evoked by acoustic stimulation and the duration of wild running and clonic seizures in the group with 20mg/kg Atomoxetine and the group with 5mg/kg Atomoxetine +100mg/kg 5-HTP were significantly reduced compared to the group with Saline. The duration of tonic-clonic seizures in the group with 10mg/kg Atomoxetine, 15mg/kg Atomoxetine, 20mg/kg Atomoxetine and the group with 5mg/kg Atomoxetine +100mg/kg 5-HTP was significantly reduced compared to the group with Saline. The seizure score in the group with 5mg/kg Atomoxetine +100mg/kg 5-HTP was markedly lower than the group with Saline. No obvious differences were observed between treatment groups in the AGSz latency (*p* > 0.05). **(E)** Schematic illustration of the observations in the DBA/1 mice induced by PTZ injection following IP injection of 5-HTP with or without Atomoxetine. **(F)** Schematic illustration of the observations in the DBA/1 mice induced by PTZ injection following IP injection of Atomoxetine with or without 5-HTP. **(G1-5)** The incidence of S-IRA evoked by PTZ, the duration of wild running and clonic seizures, the duration of tonic-clonic seizures and the seizure score in the group with 125mg/kg 5-HTP and the group with 100mg/kg 5-HTP + 5mg/kg Atomoxetine were significantly reduced compared to the group with Saline. The GSz latency in the group with 100mg/kg 5-HTP + 5mg/kg Atomoxetine was markedly higher than the group with Saline. **(H1-5)** The incidence of S-IRA evoked by PTZ, the duration of wild running and clonic seizures, the duration of tonic-clonic seizures and the seizure score in the group with 15mg/kg Atomoxetine and the group with 5mg/kg Atomoxetine + 100mg/kg 5-HTP were significantly reduced compared to the group with Saline. The GSz latency in the group with 5mg/kg Atomoxetine + 100mg/kg 5-HTP was markedly higher than the group with Saline. ****P < 0.0001; ***P < 0.001; **P < 0.01; *P < 0.05; Data are mean ± SEM; i.p.: intraperitoneal injection

### 3.2 DR-LC neural pathways synergistically regulated the occurrence of SUDEP in peripheral and central dual verification

Our experimental findings reveal that elevating 5-HT and NE neurotransmitter levels in the brain reduces S-IRA incidence in a dose-dependent manner. Combining these neurotransmitters in small doses yields a cascade amplification effect. Drawing on numerous studies and prior experiments, we identified the dorsal raphe nucleus (DR) and the locus coeruleus (LC) as the respective synthesis sites for 5-HT and NE in the brain. Thus, we selected DR and LC as target nuclei to test our hypothesis. Building on past work, we utilized venlafaxine, a selective serotonin and norepinephrine reuptake inhibitor (SNRIs), as a strategic tool to hinder the reuptake of 5-HT and NE neurotransmitters, thereby enhancing their accessibility within neuronal synapses. This approach aims to optimize the availability of monoamine transmitters, further reinforcing our understanding of their role in mitigating S-IRA.

Thirty min prior to the application of sound stimulation, different doses of venlafaxine or saline were administered to already modeled DBA/1 mice by IP injection (Fig. 2A), and the results showed that venlafaxine significantly reduced the incidence of S-IRA in a dose-dependent manner compared to the control group (P<0.0001, Fig. 2 C1). Again, to further mitigate the potential differences generated by the S-IRA induction method alone, we employed a PTZ-induced epilepsy model to validate this phenomenon. Compared with the control group, the incidence of S-IRA was significantly reduced when venlafaxine was injected intraperitoneally at 25 mg/kg (P<0.05, Fig. 2 D1), while there was no significant difference between the control group and other venlafaxine groups (P> 0.05, Fig.2 D1). The above results suggest that, unlike the acoustic stimulation model, the incidence of S-IRA, the duration of tonic-clonic seizures and the seizure score in the PTZ model decrease first and then increase with the increase of venlafaxine dose, which may be related to the different induction methods of SUDEP in the two models. Based on the results of the above two stimulation models, we have clarified the synergistic inhibitory effect of DR-LC nuclei on S-IRA, but the specific receptors need to be further clarified.

**Figure 2.**
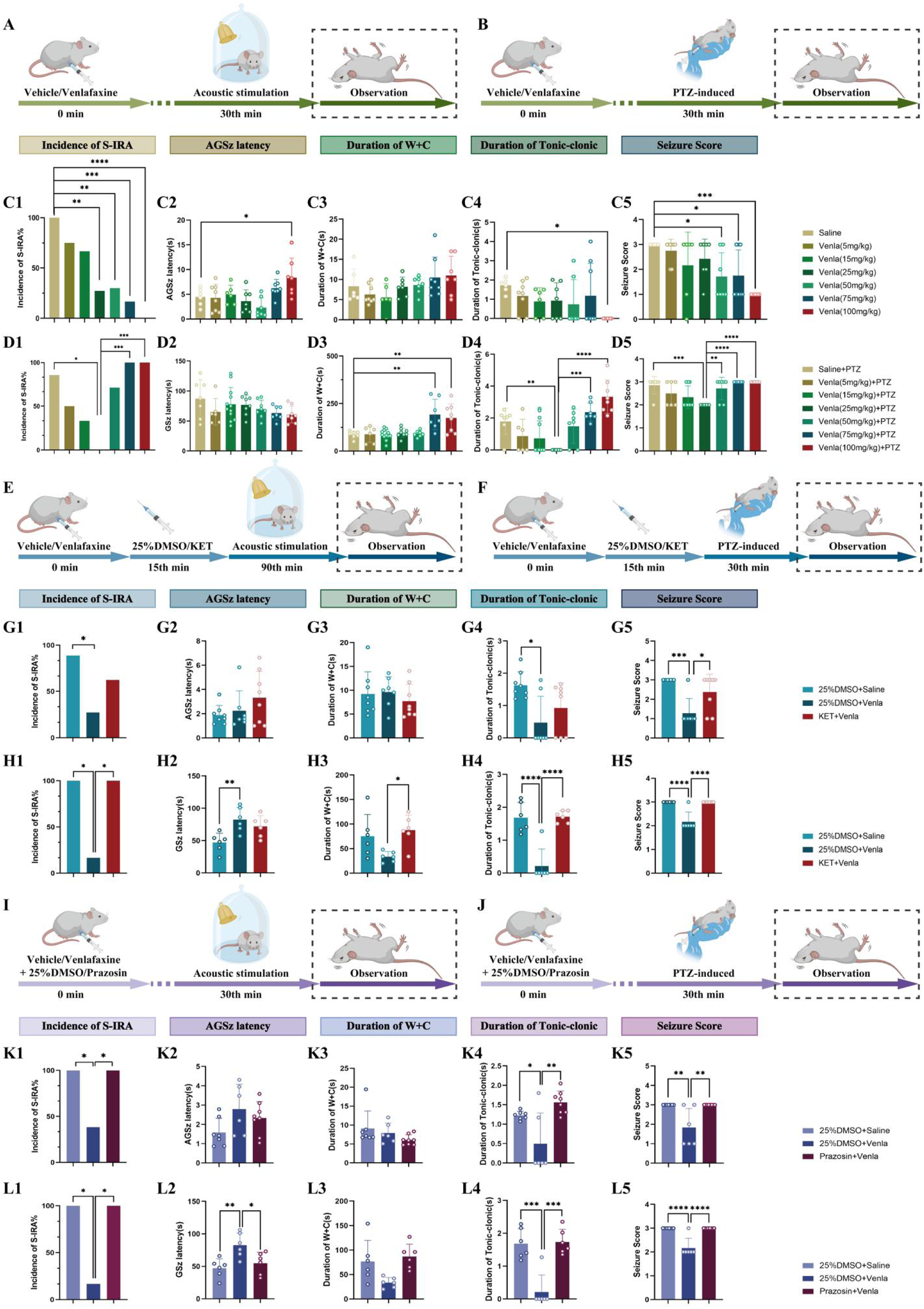
Venlafaxine significantly reduced the incidence of S-IRA induced by acoustic stimulation and PTZ injection, and 5-HT2A and NE-α1 receptor antagonists can reverse the effects. **(A)** Schematic illustration of the observations in the DBA/1 mice induced by acoustic stimulation following IP injection of different doses of Venlafaxine. **(B)** Schematic illustration of the observations in the DBA/1 mice induced by PTZ injection following IP injection of different doses of Venlafaxine. **(C1-5)** The incidence of S-IRA evoked by acoustic stimulation and the seizure score in the group with 25mg/kg Venlafaxine, 50mg/kg Venlafaxine, 75mg/kg Venlafaxine and 100mg/kg Venlafaxine were significantly reduced compared to the group with Saline. The AGSz latency in the group with 100mg/kg Venlafaxine was markedly higher than the group with Saline. The duration of tonic-clonic seizures in the group with 100mg/kg Venlafaxine was markedly higher than the group with Saline. The seizure score in the group with 50mg/kg Venlafaxine, 75mg/kg Venlafaxine and 100mg/kg Venlafaxine was significantly decreased compared to the group with Saline. No obvious differences were observed between treatment groups in the duration of wild running and clonic seizures (*p* > 0.05). **(D1-5)** The incidence of S-IRA evoked by PTZ and the duration of tonic-clonic seizures in the group with 25mg/kg Venlafaxine were markedly reduced compared to the group with Saline and the group with 75mg/kg Venlafaxine and 100mg/kg Venlafaxine. The duration of wild running and clonic seizures in the group with 75mg/kg Venlafaxine and 100mg/kg Venlafaxine was markedly higher than the group with Saline. The seizure score in the group with 25mg/kg Venlafaxine was significantly decreased compared to the group with Saline and the group with Saline, 50mg/kg Venlafaxine, 75mg/kg Venlafaxine and 100mg/kg Venlafaxine. No obvious differences were observed between treatment groups in the GSz latency (*p* > 0.05). **(E)** Schematic illustration of the observations in the DBA/1 mice induced by acoustic stimulation following IP injection of Venlafaxine and 5-HT2A receptor antagonist KET. **(F)** Schematic illustration of the observations in the DBA/1 mice induced by PTZ injection following IP injection of Venlafaxine and 5-HT2A receptor antagonist KET. **(G1-5)** The incidence of S-IRA evoked by acoustic stimulation, the duration of tonic-clonic seizures and the seizure score in the group with 25%DMSO + Venlafaxine were markedly lower compared with other groups. No obvious differences were observed between treatment groups in the AGSz latency and the duration of wild running and clonic seizures (*p* > 0.05). **(H1-5)** The incidence of S-IRA evoked by PTZ injection, the duration of tonic-clonic seizures and the seizure score in the group with 25%DMSO + Venlafaxine were markedly lower compared with other groups. The GSz latency in the group with 25%DMSO + Venlafaxine was significantly increased compared to the group with 25%DMSO + Saline. The duration of wild running and clonic seizures in the group with 25%DMSO + Venlafaxine was markedly decreased compared to the group with KET + Venlafaxine. **(I)** Schematic illustration of the observations in the DBA/1 mice induced by acoustic stimulation following IP injection of Venlafaxine and NE-α1 receptor antagonist Prazosin. **(J)** Schematic illustration of the observations in the DBA/1 mice induced by PTZ injection following IP injection of Venlafaxine and NE-α1 receptor antagonist Prazosin. **(K1-5)** The incidence of S-IRA evoked by acoustic stimulation, the duration of tonic-clonic seizures and the seizure score in the group with 25%DMSO + Venlafaxine were lower compared with other groups. No obvious differences were observed between treatment groups in the AGSz latency and the duration of wild running and clonic seizures (*p* > 0.05). **(L1-5)** The incidence of S-IRA evoked by PTZ injection, the duration of tonic-clonic seizures and the seizure score in the group with 25%DMSO + Venlafaxine were markedly lower compared with other groups. The GSz latency in the group with 25%DMSO + Venlafaxine was markedly increased compared with other groups. No obvious differences were observed between treatment groups in the duration of wild running and clonic seizures (*p* > 0.05). ****P < 0.0001; ***P < 0.001; **P < 0.01; *P < 0.05; Data are mean ± SEM; i.p.: intraperitoneal injection

To investigate whether Venlafaxine acts on the receptor after previous studies have shown that 5-HT2A receptor and NE-α1 receptor are closely related to the regulation of respiratory function, we injected the 5-HT2A receptor antagonist KET and the NE-α1 receptor antagonist Prazosin IP-injected to observe the behavioral changes of SUDEP in mice (Fig.2 E-F, I-J). In two models of acoustic stimulation nuclear PTZ injection, the findings revealed that subsequent administration of KET subsequent to Venlafaxine’s IP injection, in contrast to the 25% DMSO + Venlafaxine 25 mg/kg group, not only negated Venlafaxine’s efficacy in decreasing the prevalence of S-IRA but also reinstated higher epilepsy scores (Fig.2 G1-5, H1-5). Furthermore, when comparing with the same control group, the IP injection of a combination of Prazosin and Venlafaxine markedly reversed Venlafaxine’s suppressive effect on S-IRA incidence, notably extending the duration of tonic-clonic seizures and escalating seizure severity scores (Fig.2 K1-5, L1-5).

Based on Venlafaxine’s pharmacological properties, where it acts solely on 5-HT receptors at low doses, similar to SSRIs, and on both 5-HT and NE receptors at medium to high doses, we aimed to specifically investigate whether Venlafaxine, at its minimum effective dose for reducing SUDEP incidence, exerts its effects concurrently through both the 5-HT and NE systems. To achieve this, we utilized PCPA (an inhibitor of TPH, the rate-limiting enzyme in 5-HT synthesis) and DSP-4 (a selective neurotoxin for noradrenergic neurons) for our observation and analysis (Fig.3 A, C). Compared with the experimental group, the incidence of S-IRA was significantly increased after 5 days of pretreatment with PCPA with venlafaxine injection compared with the experimental group with venlafaxine alone, reversing the reduction of S-IRA by Venlafaxine (P<0.01, Fig.3B). Similarly, the incidence of S-IRA in the IP injection of Venlafaxine 1 day after DSP-4 pretreatment was significantly higher than that in the Venlafaxine alone group, reversing the phenomenon of Venlafaxine reducing the occurrence of S-IRA (P< 0.05, Fig.3D). We were also pleasantly surprised to find that PCPA pretreatment 5 days + DSP-4 pretreatment 1 day were able to completely reverse the S-IRA reduction with Venlafaxine (P < 0.0001, Fig.3B5).

**Figure 3.**
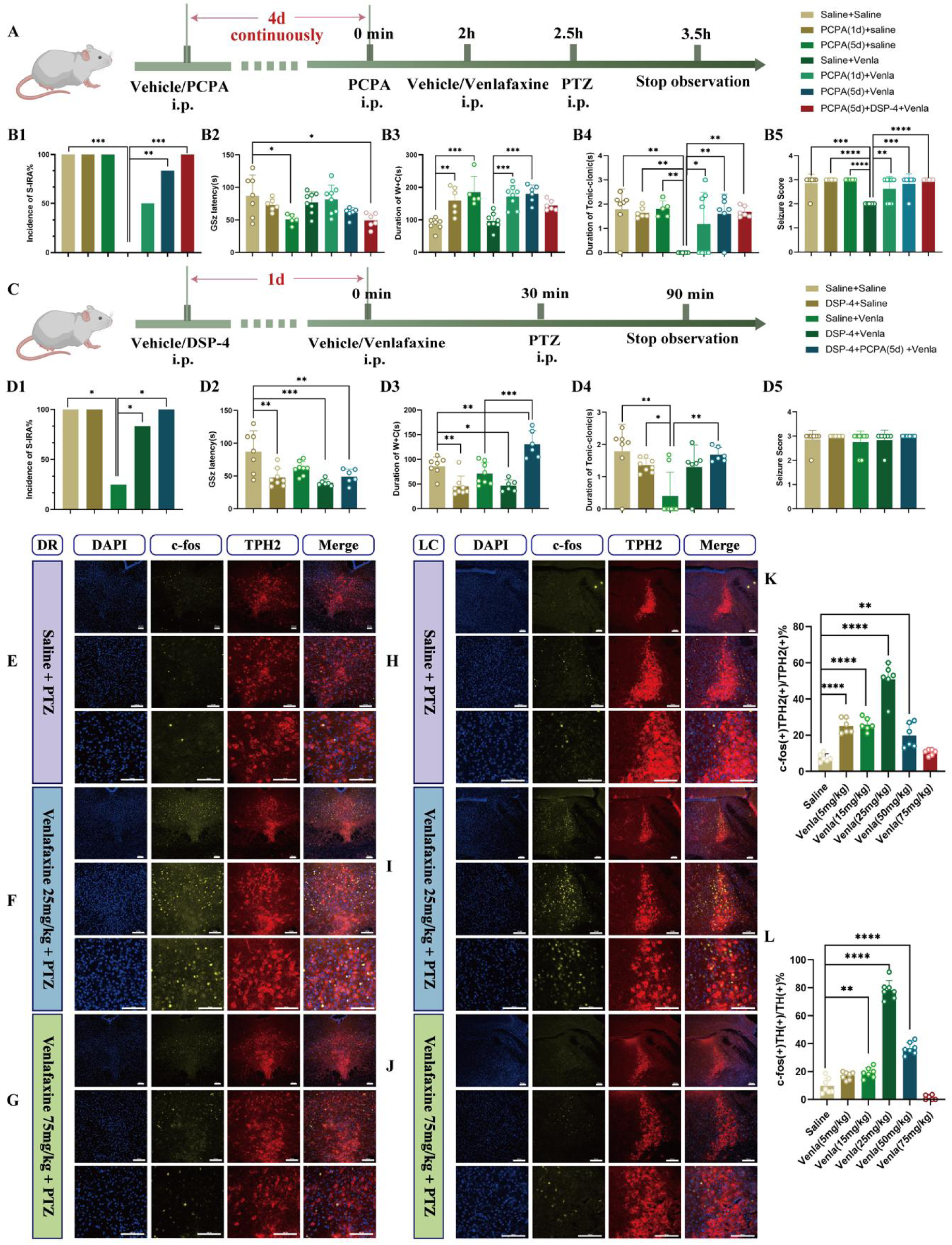
Both DR 5-HT neurons and LC NE neurons are involved in the regulation of SUDEP. **(A)** Experimental protocol for the observations in the DBA/1 mice induced by PTZ injection following IP injection of PCPA and Venlafaxine. **(B1-5)** The incidence of S-IRA evoked by PTZ in the group with Venlafaxine was significantly reduced compared to the group with Saline, the group with PCPA (5d) + Venlafaxine and the group with PCPA (5d) + DSP-4 + Venlafaxine. The GSz latency in the group with PCPA (5d) and the group with PCPA (5d) + DSP-4 + Venlafaxine was markedly lower than the group with Saline. The duration of wild running and clonic seizures in the group with PCPA (1d) and the group with PCPA (5d) was markedly higher than the group with Saline, and the same significant goes for the group with Venlafaxine, the group with PCPA (1d) + Venlafaxine and the group with PCPA (5d) + Venlafaxine. The duration of tonic-clonic seizures and seizure score in the group with Venlafaxine was significantly lower compared with the other groups. **(C)** Experimental protocol for the observations in the DBA/1 mice induced by PTZ injection following IP injection of DSP-4 and Venlafaxine. **(D1-5)** The incidence of S-IRA evoked by PTZ in the group with Venlafaxine was significantly reduced compared to the group with Saline, the group with DSP-4 + Venlafaxine and the group with DSP-4 + PCPA (5d) + Venlafaxine. The GSz latency in the group with DSP-4, the group with DSP-4 + Venlafaxine and the group with DSP-4 + PCPA (5d) + Venlafaxine was markedly lower than the group with Saline. The duration of wild running and clonic seizures in the group with DSP-4, the group with DSP-4 + Venlafaxine and the group with DSP-4 + PCPA (5d) + Venlafaxine was markedly decreased than the group with Saline. The duration of wild running and clonic seizures in the group with Venlafaxine was significantly decreased than the group with DSP-4 + PCPA (5d) + Venlafaxine. The duration of tonic-clonic seizures in the group with Venlafaxine was significantly lower compared with the group with Saline, the group with DSP-4 and the group with DSP-4 + PCPA (5d) + Venlafaxine. No obvious differences were observed between treatment groups in the seizure score (*p* > 0.05). **(E-G)** Representative images of staining for c-fos, TPH2 and DAPI in the DR in the groups with Saline + PTZ, Venlafaxine 25mg/kg + PTZ and Venlafaxine 75mg/kg + PTZ. **(H-J)** Representative images of staining for c-fos, TH and DAPI in the LC in the groups with Saline + PTZ, Venlafaxine 25mg/kg + PTZ and Venlafaxine 75mg/kg + PTZ. **(K)** The quantification of c-fos (+)/TPH2 (+) section in the DR. Significantly more c-fos (+)/TPH2 (+) cells were observed in the group with 5mg/kg Venlafaxine, 15mg/kg Venlafaxine, 25mg/kg Venlafaxine and 50mg/kg Venlafaxine. **(L)** The quantification of c-fos (+)/TH (+) section in the LC. Significantly more c-fos (+)/TH (+) cells were observed in the group with 15mg/kg Venlafaxine, 25mg/kg Venlafaxine and 50mg/kg Venlafaxine. ****P < 0.0001; ***P < 0.001; **P < 0.01; *P < 0.05; Data are mean±SEM; i.p.: intraperitoneal injection

Drawing from the aforementioned results, we conclude that Venlafaxine’s capacity to diminish S-IRA incidence, mediated via the 5-HT2A and NE-α1 receptors, can be reversed by impeding or dampening the influence of either receptor. Consequently, we posit that the DR and LC neuronal groups exhibit a synergistic, interdependent mechanism in modulating the occurrence of S-IRA. To substantiate this hypothesis, we conducted immunohistochemistry experiments, which provided further clarity and confirmation.

TPH2 is a rate-limiting enzyme for the synthesis of 5-HT, and its expression level is closely related to the activity of 5-HT synthesis, and is mainly expressed in 5-HT neurons. TH is a key enzyme in the NE synthesis pathway, which catalyzes the conversion of tyrosine to dopamine, which is the first step in the NE synthesis pathway, and the expression of TH can also reflect the activity and functional status of NE neurons, so we chose TPH2 and TH to represent 5-HT neurons within DR and NE neurons within LC, respectively (Fig. 3E-J). The results showed that in the PTZ model, the expression of c-fos in 5-HT neurons in DR and NE neurons in LC increased first and then decreased with the increase of Venlafaxine dose. Compared with the control group, the c-fos of 5-HT neurons in the DR were significantly increased at 5 mg/kg, 15 mg/kg, 25 mg/kg, and 50 mg/kg of Venlafaxine, suggesting significant activation of 5-HT neurons (P < 0.001, Fig.3K). Venlafaxine was 15 mg/kg (P < 0.01), 25 mg/kg (P < 0.001) and 50 mg/kg (P<0.001), the expression of c-fos in NE neurons in the LC was significantly increased compared to the control group, indicating that NE neurons were significantly activated (Fig.3L). These results suggest that both 5-HT neurons in DR and NE neurons in LC are involved in the mechanism of regulating the occurrence of SUDEP, and with the increase of Venlafaxine dose, 5-HT neurons in DR and NE neurons in LC may be activated first and then inhibited, which may be related to positive and negative feedback caused by drugs.

Based on the above experimental results, we have verified the role of DR5-HT and LC-NE neurons in the regulation of S-IRA. So far, we have finally confirmed the inhibition of SUDEP by the DR-LC neural pathway, especially from the peripheral pathway. However, considering the lag of c-fos expression, we will further employ real-time monitoring methods to determine the expression neurons activity of the DR-LC neural pathway

### 3.3 Calcium signaling results validated the regulation of SUDEP by the DR-LC neural pathway at the prominent level

To further determine the specificity target nuclei and locations within the central nervous system, we implanted cannulas in the lateral ventricles of mice and injected specific calcium signaling viruses into the DR and LC, respectively. Three weeks after infection, we conducted calcium signaling recording experiments. Ten minutes after the calcium signaling recording, different doses of Venlafaxine (2.0ul) were administered through the ICV cannulas. Ten minutes later, PTZ was injected intraperitoneally, and changes in the activity of 5-HT neurons in the DR and NE neurons in the LC were recorded (Fig.4 A-C). The results showed that compared with the control group, the incidence of S-IRA in mice was significantly reduced when Venlafaxine was administered at 6.25mg/ml (P<0.05, Fig.4 E). However, as the concentration of Venlafaxine increased, the incidence of S-IRA in mice gradually increased from low to high. When Venlafaxine was administered at 25mg/ml, the incidence of S-IRA was significantly higher than that in the 6.25mg/ml group (P<0.05, Fig.4 E). This trend is consistent with the trend of S-IRA incidence in the PTZ model when different concentrations of Venlafaxine were injected intraperitoneally. During clonic seizures in mice, compared with the control group, the calcium signaling changes of 5-HT neurons in the DR began to increase at Venlafaxine 1.25mg/ml (P <0.01, Fig.4 I). As the concentration of Venlafaxine increased, the calcium signaling changes showed a trend of first increasing and then decreasing (Fig.4 I). Similarly, we also observed that during clonic seizures, compared with the control group, the calcium signaling changes of NE neurons in the LC began to increase significantly at Venlafaxine 2.5mg/ml and peaked at Venlafaxine 6.25mg/ml (P <0.001, Fig.4 I). As the concentration of Venlafaxine increased, the calcium signaling changes also showed a trend of first increasing and then decreasing (Fig.4 I).

**Figure 4.**
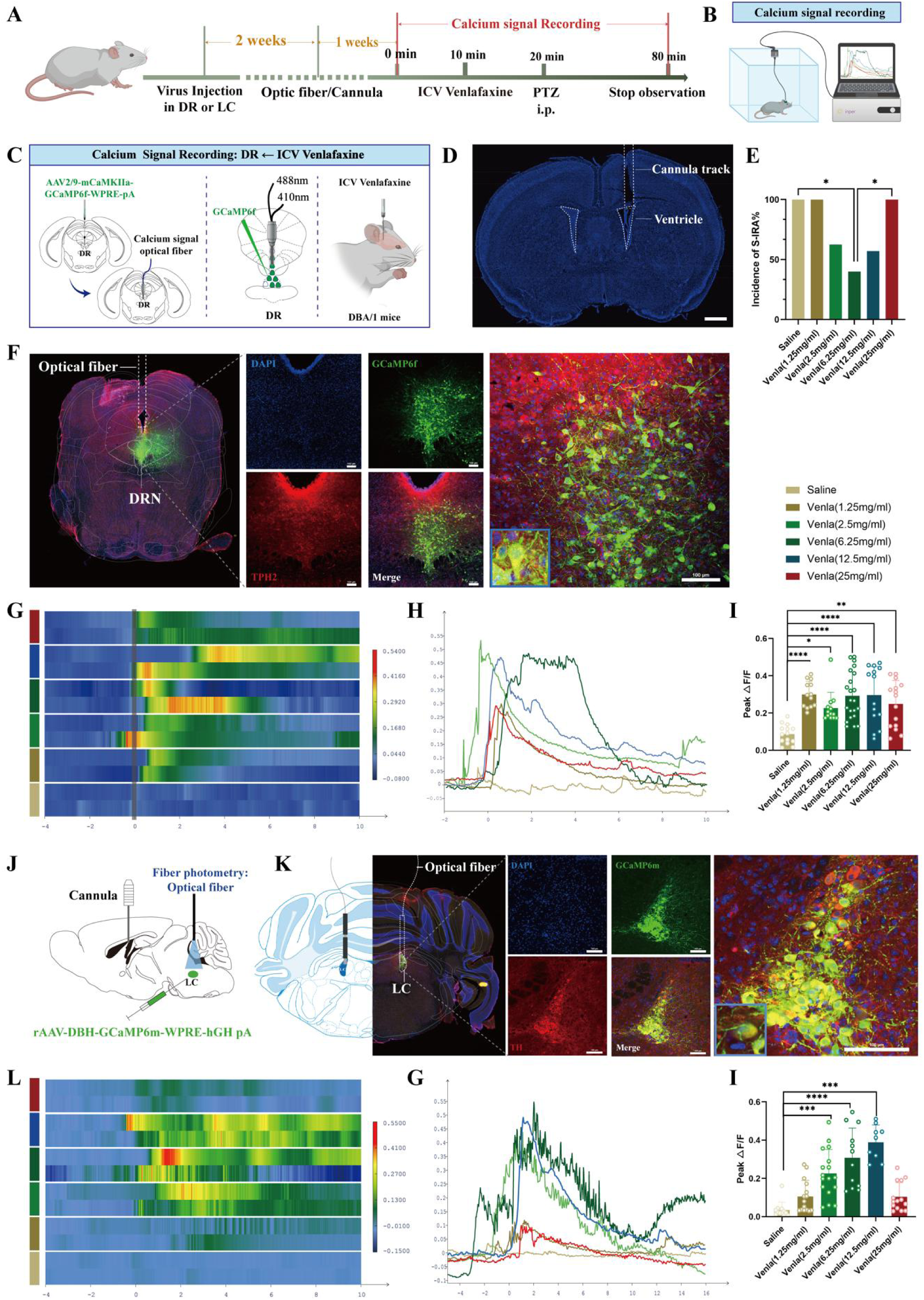
Calcium signaling records tonic seizures in DR 5-HT neuron and LC NE neuron demonstrate the effect of reducing the incidence of S-IRA by activation of both neurons. **(A)** Experimental protocol for calcium signal recording of DR 5-HT neurons or LC NE neurons and ICV injection of Venlafaxine in the PTZ-induced TH-Cre DBA/1 mice. **(B)** A schematic of the calcium signal recording device. **(C)** Schematic diagram of the positions for virus injection and optic fibers implantation in the DR and ICV injection of Venlafaxine. **(D)** Representative image of the track of cannula implanted for ICV injection. **(E)** The incidence of S-IRA evoked by PTZ in the group with 6.25mg/ml Venlafaxine was markedly reduced compared to the group with Saline and the group with 25mg/ml Venlafaxine. **(F)** Representative images show the optical fiber location and the co-expression of GCaMP6f, TPH2 and DAPI in the DR. **(G-H)** Heatmap and statistical diagram of calcium signaling changes in DR during the observation phase from two representative mice in the groups with ICV injection of Saline or different dose of Venlafaxine. **(I)** The peak ΔF/F in DR of tonic seizures was significantly increased in all groups with ICV injection of different dose of Venlafaxine than in the control group. **(J)** Schematic diagram of the positions for virus injection and optic fibers implantation in the LC and ICV injection of Venlafaxine. **(K)** Representative images show the optical fiber location and the co-expression of GCaMP6f, TH and DAPI in the LC. **(L-M)** Heatmap and statistical diagram of calcium signaling changes in LC during the observation phase from two representative mice in the groups with ICV injection of Saline or different dose of Venlafaxine. **(N)** The peak ΔF/F in DR of tonic seizures was significantly increased in the groups with ICV injection of 2.5mg/ml Venlafaxine, 6.25mg/ml Venlafaxine and 12.5mg/ml Venlafaxine than in the control group. ****P < 0.0001; ***P < 0.001; **P < 0.01; *P < 0.05; Data are mean±SEM; i.p.: intraperitoneal injection; icv: intracerebroventricular

These results further suggest that, similar to peripheral administration, we applied the expression of calcium signal at the synaptic level to verify that after intraventricular administration, 5-HT neurons in DR and NE neurons in LC were first activated and then inhibited with the increase of venlafaxine concentration, which was consistent with the statistical trend of the incidence of S-IRA in mice, and once again provided strong evidence for the regulation of S-IRA by DR^5-HT^-LC^NE^ nerves through synergy dependence from the perspective of central regulation.

### 3.4 Specific stimulation of the DR^5-HT^ and LC^NE^ nuclei revealed the sequential order of the DR-LC neural pathway

Despite validating our conclusions on the DR-LC neural pathway’s synergistic regulation of S-IRA incidence through rigorous peripheral pharmacological approaches and central calcium signaling analysis, there remains a gap in specific validation tailored specifically to this finding. Given the intimate interplay between these two entities, we endeavor to elucidate their mutual regulatory dynamics and delve into the intricate upstream and downstream sequences that govern the function of the DR^5-HT^ and LC^NE^, thereby enhancing the comprehensiveness of our understanding.

To investigate the specific roles of 5-HT neurons in DR and NE neurons in LC in SUDEP and the connection between them, we injected optogenetic viruses into DR and performed optogenetic experiments three weeks after infection. For the DR light-stimulated group: 15mv20min, with PTZ injected intraperitoneally at the 15th minute of light stimulation. The behavior of mice was observed and recorded, and the c-fos expression of NE neurons in LC of mice was statistically analyzed by immunohistochemistry (Fig.5 A). The results showed that after activating 5-HT neurons in DR with light stimulation, compared with the non-light-stimulated group, the incidence of S-IRA in the light-stimulated group was significantly reduced, the latency of epilepsy was significantly prolonged, the duration of tonic spasms was shortened, and the epilepsy score was decreased (P<0.05, Fig.5 E1-E5). Meanwhile, it was also found that compared with the non-light-stimulated group, NE neurons in LC of the light-stimulated group were significantly activated, and the expression of c-fos in NE neurons in the left and right LC was significantly increased (P<0.001, Fig.5 F). This indicates that activating 5-HT neurons in DR can significantly reduce the incidence of S-IRA and also significantly activate NE neurons in LC.

**Figure 5.**
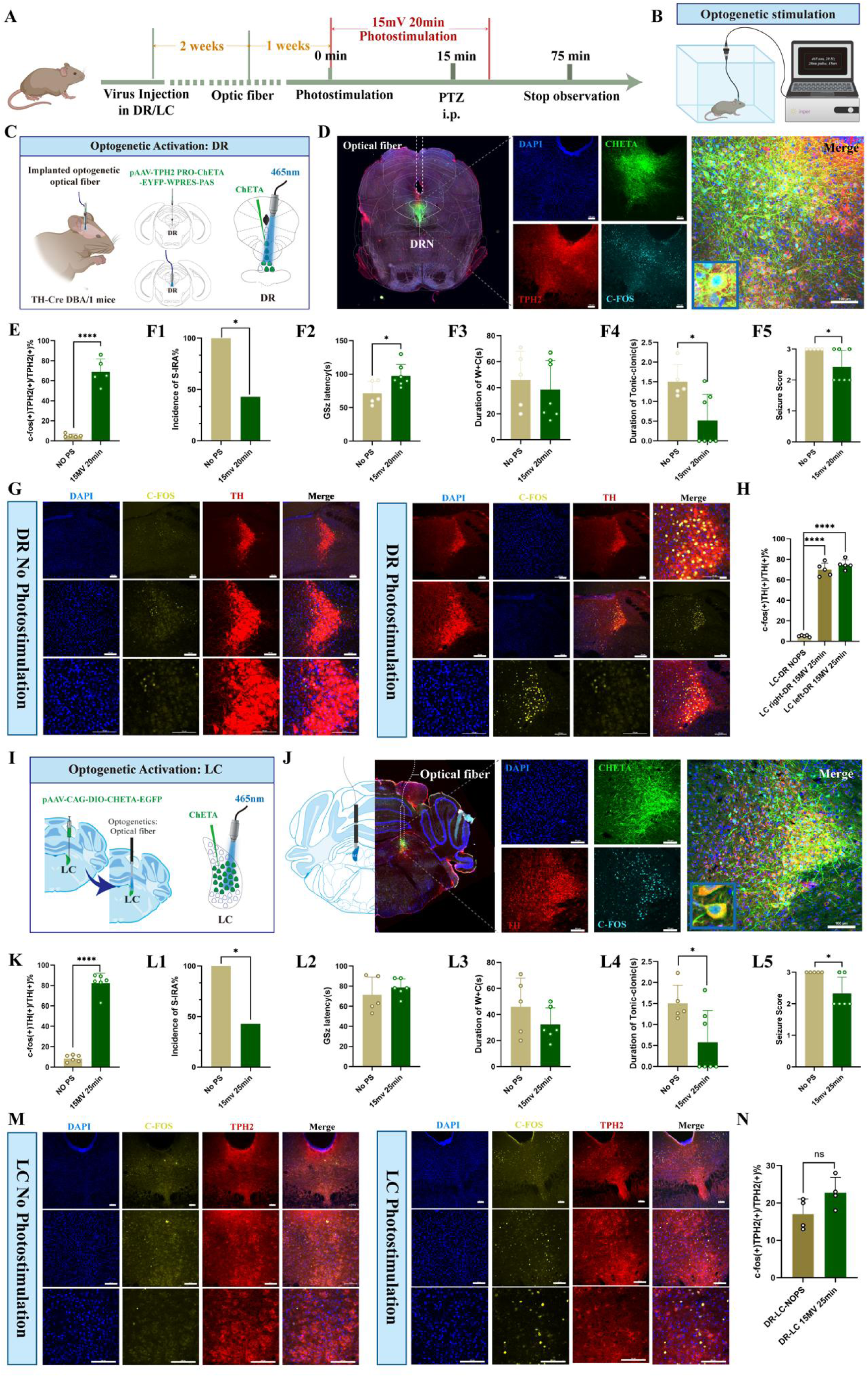
Optogenetic activation of DR 5-HT neurons and LC NE neurons reduced the incidence of S-IRA evoked by PTZ. **(A)** Experimental protocol for optogenetic stimulation of DR 5-HT neurons or LC NE neurons and ICV injection of Venlafaxine in the PTZ-induced TH-Cre DBA/1 mice. **(B)** A schematic of the optogenetic stimulation device. **(C)** Schematic diagram of the positions for virus injection and optic fibers implantation in the DR. **(D)** Representative images show the optical fiber location and the co-expression of CHETA, TPH2, C-FOS and DAPI in the DR. **(E)** The quantification of c-fos (+)/TPH2 (+) section in the DR with or without DR photostimulation Significantly more c-fos (+)/TPH2 (+) cells in the DR were observed in the group with photostimulation (15mv 20min). **(F1-5)** The incidence of S-IRA evoked by PTZ injection, the duration of tonic-clonic seizures and the seizure score in the group with DR photostimulation (15mv 20min) were markedly reduced compared to the group without DR photostimulation. The GSz latency in the group with DR photostimulation (15mv 20min) were markedly increased compared to the group without DR photostimulation. No obvious differences were observed between treatment groups in the duration of wild running and clonic seizures (*p* > 0.05). **(G)** Representative images of staining for C-FOS, TH, and DAPI in the LC in the groups with and without DR photostimulation. **(H)** The quantification of c-fos (+)/TH (+) section in the LC with or without DR photostimulation. Significantly more c-fos (+)/TH (+) cells in the bilateral LC were observed in the group with DR photostimulation. **(I)** Schematic diagram of the positions for virus injection and optic fibers implantation in the LC. **(J)** Representative images show the optical fiber location and the co-expression of CHETA, TH, C-FOS and DAPI in the LC. **(K)** The quantification of c-fos (+)/TH (+) section in the LC with or without LC photostimulation. Significantly more c-fos (+)/TH (+) cells were observed in the group with photostimulation (15mv 25min). **(L1-5)** The incidence of S-IRA evoked by PTZ injection, the duration of tonic-clonic seizures and the seizure score in the group with LC photostimulation (15mv 20min) were markedly reduced compared to the group without LC photostimulation. No obvious differences were observed between treatment groups in the GSz latency and the duration of wild running and clonic seizures (*p* > 0.05). **(M)** Representative images of staining for C-FOS, TPH2, and DAPI in the DR in the groups with and without LC photostimulation. **(N)** The quantification of c-fos (+)/TPH2 (+) section in the DR with or without LC photostimulation. Significantly more c-fos (+)/TPH2 (+) cells were observed in the group with LC photostimulation. ****P < 0.0001; *P < 0.05; ns: P > 0.05; Data are mean±SEM; PS: photostimulation; NO PS: no photostimulation; i.p.: intraperitoneal injection;

Meanwhile, we injected optogenetic viruses into the bilateral LC of TH-cre transgenic DBA/1 background mice. After 3 weeks of infection, we conducted optogenetic experiments. For the bilateral LC light stimulation group: 15mv for 25min, and PTZ was injected intraperitoneally at the 20th minute of light stimulation. We observed and recorded the behavior of mice and immunohistochemically counted the c-fos expression of 5-HT neurons in the DR of mice (Fig.5 A). The results showed that light stimulation activated NE neurons in the bilateral LC. Compared with the non-light stimulation group, the incidence of S-IRA in the light stimulation group was significantly reduced, and the duration of tonic convulsion was also significantly shortened, resulting in a lower epilepsy score (P<0.05, Fig.5 J1, J4-J5). However, there were no significant differences in the latency of GSz onset and the duration of running convulsion (P>0.05, Fig.5 J2-J3). It was also found that compared with the non-light stimulation group, there was no significant difference in the expression of c-fos in 5-HT neurons in the DR of the bilateral LC light stimulation group (P>0.05, Fig.5 K). This indicated that optogenetic activation of NE neurons in the bilateral LC could significantly reduce the incidence of S-IRA but could not significantly activate 5-HT neurons in the DR.

To further investigate the specific roles of 5-HT neurons in DR and NE neurons in LC in SUDEP and the connection between them, we injected chemogenetic viruses into the bilateral LC of TH-cre transgenic DBA/1 background mice. Following three weeks of infection, chemogenetic experiments were conducted. Twenty minutes later, different doses of CNO were administered intraperitoneally, followed by PTZ injection 30 minutes later. Mouse behavior was observed and recorded, and immunohistochemical analysis was performed to quantify c-fos expression in 5-HT neurons within the DR of mice (Fig.6 A). The results showed that chemogenetic activation of NE neurons in the bilateral LC significantly reduced the incidence of S-IRA in the 1mg/kg CNO group compared to the saline group, while there was no significant difference in the 0.5mg/kg CNO group compared to the saline group (P<0.05, Fig.6 D1).In addition, we found that compared with the control group, the latency of epilepsy was prolonged, the duration of tonic convulsion was shortened, and the epilepsy score was decreased in the group with 1mg/kg CNO injected intraperitoneally. Furthermore, compared to the control group, the expression of c-fos in 5-HT neurons within the DR was significantly increased in the 1mg/kg CNO group with bilateral LC stimulation (P>0.05, Fig.6 F). This suggests that chemogenetic activation of NE neurons in the bilateral LC at a certain intensity can significantly reduce the incidence of S-IRA while also markedly activating 5-HT neurons in the DR.

**Figure 6.**
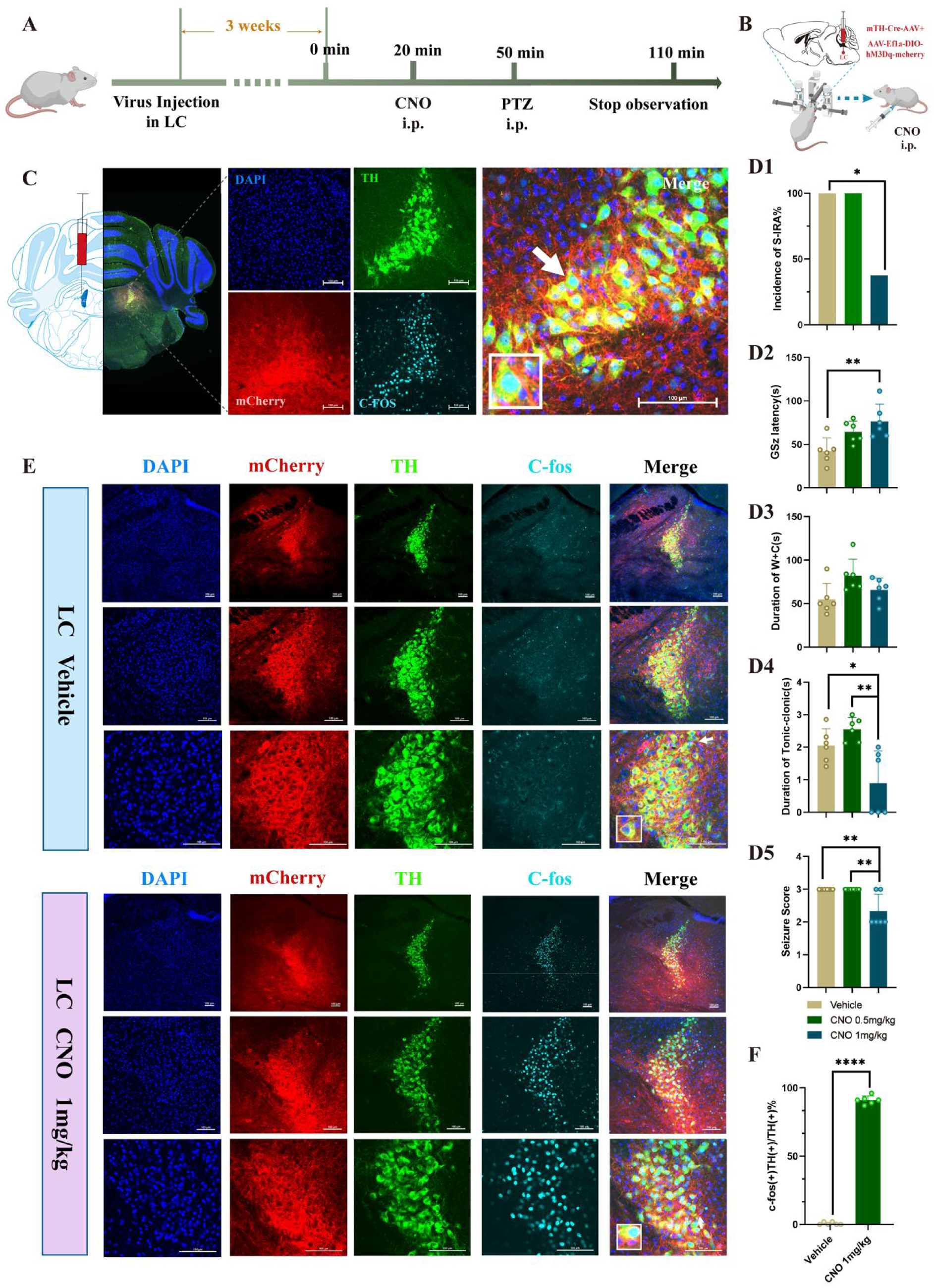
Chemogenetic activation of LC NE neurons significantly reduced the incidence of S-IRA evoked by PTZ. **(A)** Experimental protocol for chemogenetic activation of LC NE neurons in the PTZ-induced DBA/1 mice. **(B)** Schematic diagram of the location of virus (mTH-Cre-AAV+AAV-EF1a-DIO-hM3Dq-mCherry) injection in LC and IP injection of CNO. **(C)** Representative images show the location of virus injection and the co-expression of TH, mCherry, C-FOS and DAPI in the LC. **(D1-5)** The incidence of S-IRA evoked by PTZ in the group with 1mg/kg CNO was markedly reduced compared to the group with Vehicle. The GSz latency in the group with 1mg/kg CNO was significantly higher than the group with Vehicle. The duration of tonic-clonic seizures and the seizure score in the group with 1mg/kg CNO was significantly lower than the group with Vehicle and the group with 0.5mg/kg CNO. No obvious differences were observed between treatment groups in the duration of wild running and clonic seizures (*p* > 0.05). **(E)** Representative images show the co-expression and staining for TH, mCherry, C-FOS and DAPI in the LC with or without CNO activation. **(F)** The quantification of c-fos (+)/TH (+) section in the LC with or without CNO activation. Significantly more c-fos (+)/TH (+) cells were observed in the 1mg/kg CNO group than in the Vehicle group (*p* < 0.0001). ****P < 0.0001; **P < 0.01; *P < 0.05; Data are mean±SEM; i.p.: intraperitoneal injection

### 3.5 Exploring the Projection Relationship of the DR-LC-PBC Neural Pathway in a Bidirectional manner across Nuclei

The above results suggest that optogenetic activation of 5-HT neurons within DR can significantly reduce the incidence of S-IRA and activate NE neurons within LC. Similarly, photo- and chemogenetic activation of NE neurons within LC can also significantly reduce the incidence of S-IRA, but cannot activate 5-HT neurons within DR. Since the two nuclei are only the production sites for the synthesis of monoaminergic neurotransmitters and do not directly regulate respiration, we combined the results of our previous research that activated 5-HT neurons in the DR can reduce the incidence of S-IRA through the 5-HT2A receptor on PBC. Since neurons within PBCs are closely related to the generation and maintenance of respiratory rhythms, although they do not project directly to phrenic motor neurons, they can coordinate respiratory activity by interacting with neurons in other brainstem regions. LC projects to premotor respiratory neurons and may affect respiratory rhythms by modulating these neurons. Therefore, we hypothesize that there is a projection relationship between DR, LC, and PBC, and that 5-HT neurons within DR may act as upstream regulators of NE neurons within LC, further regulating PBC activity through NE neurons. To test this hypothesis, we injected HSV virus into the DR for anterograde tracing to see if NE neurons within LC and neurons within PBC were labeled. To test this hypothesis, we injected HSV virus into the DR for anterograde tracing to see if NE neurons within LC and neurons within PBC were labeled. Initially, we injected the viral HSV-1 H129 into the DR for an infectious period of 36-48 hours, followed by harvesting brains for immunohistochemical analysis. (Fig. 7 A-C) showed that both sides of LC immunofluorescence staining (n=5) had yellow fluorescent markings, i.e., HSV and TH immunofluorescence co-localization expression (Fig. 7 D-E). Similarly, the same colocalization expression was observed within the PBC nucleus (Figure 7 G-H). Based on the above immunofluorescence results, we have identified the presence of neural projections between DR, LC, and PBC, and that 5-HT neurons within DR may act as upstream regulators of NE neurons within LC, further regulating PBC activity through NE neurons.

**Figure 7.**
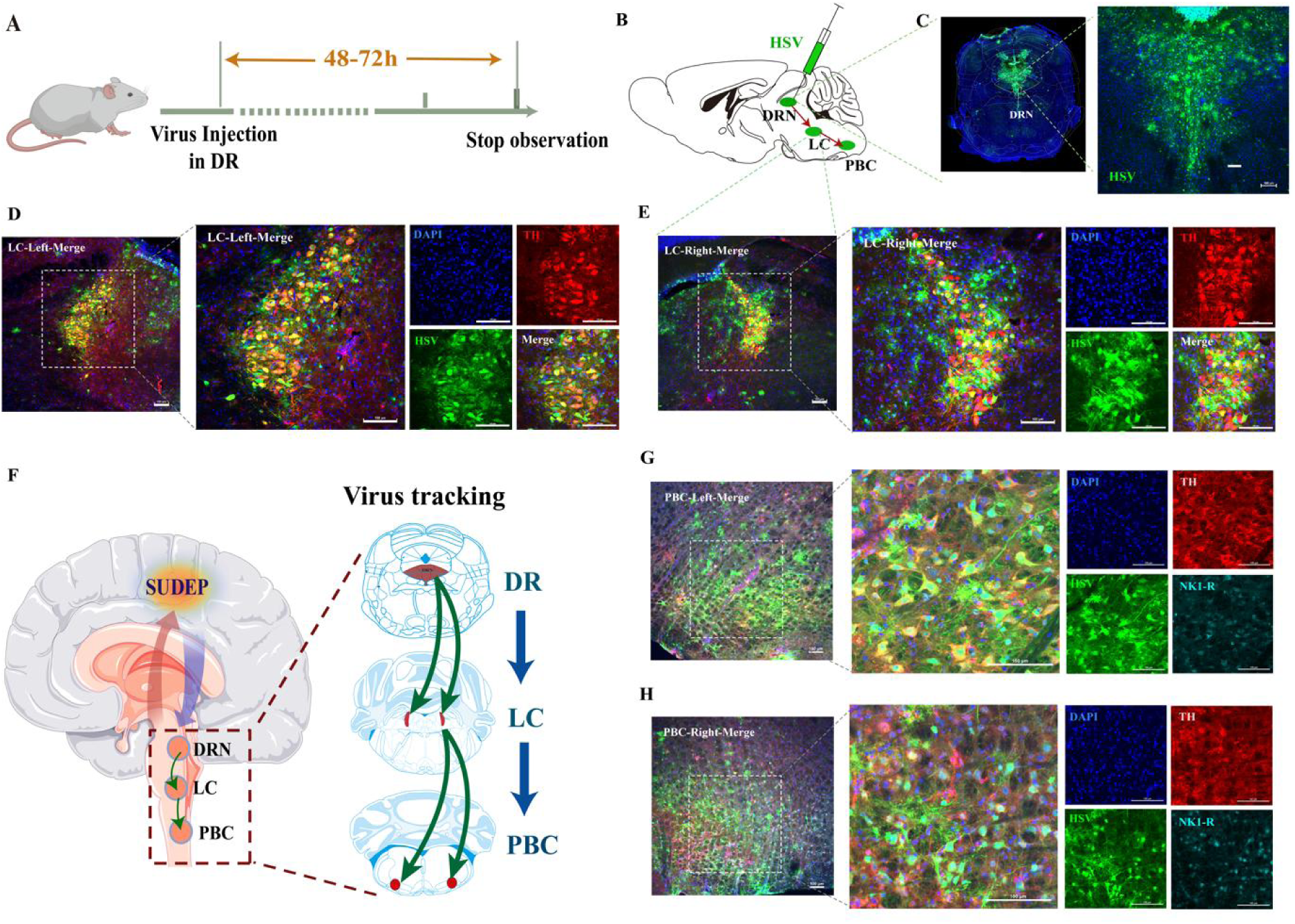
The existence of projection relationship of DR→LC→PBC neural pathway by anterograde tracing. **(A)** Experimental protocol for the anterograde tracing by injecting anterograde tracing virus HSV into DR. **(B)** Schematic of the position of the virus HSV injection and anterograde labeling. **(C)** Representative coronal brain slice, showing the location of HSV injected in the DR. **(D-E)** Representative images show the co-expression and staining of HSV, TH and DAPI in the bilateral LC. **(F)** Schematic diagram of the positions for virus and virus tracking of DR→LC→PBC neural circuit. **(G-H)** Representative images show the co-expression and staining of HSV, TH, NK1-R and DAPI in the bilateral PBC.

In addition to using the above-mentioned anterograde virus tracing method, in order to perform cis-reverse bidirectional validation, we injected CTB virus into PBC for retrograde tracing, and observed whether NE neurons in LC and 5-HT neurons in DR were labeled, further verifying that NE neurons in LC are upstream of neurons in PBC, and 5-HT neurons in DR are upstream of NE neurons in LC. Initially, the CTB virus is injected into the PBC for an infectious period of 1-2 weeks, followed by brain extraction for immunohistochemical analysis (Fig. 8 A-B). CTB was significantly labeled in DR and LC, and the fluorescence expression increased significantly (Fig.8 C-D). These results reaffirm the neural projection relationships among the DR, LC, and PBC, suggesting that 5-HT neurons in the DR are upstream of NE neurons in the LC, which in turn regulate PBC activity through NE neurons. Furthermore, 5-HT neurons in the DR may also activate neurons in the PBC by activating NE neurons in the LC, thereby reducing the occurrence of S-IRA.

**Figure 8.**
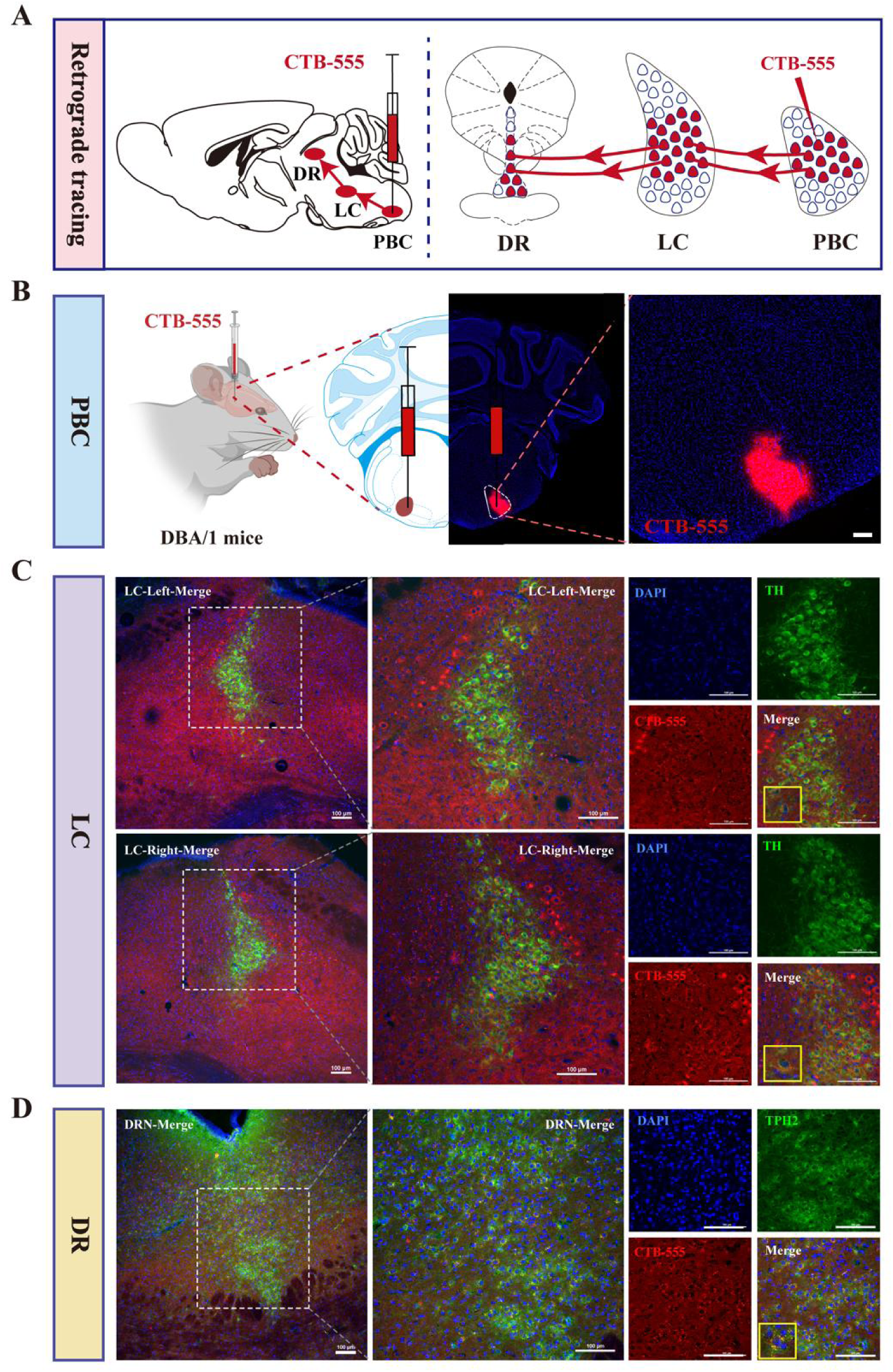
Elucidating the neural projection relationship of DR→LC→PBC neural pathway by injecting a retrograde tracer CTB-555 into PBC for dual confirmation. **(A)** Schematic of the position of the virus injection and retrograde labeling. **(B)** Representative coronal brain slice, showing the location of CTB-555 injected in the PBC. **(C)** Representative images show the co-expression and staining of CTB-555, TH and DAPI in the bilateral LC. **(D)** Representative images show the co-expression and staining of CTB-555, TPH2 and DAPI in the DR.

### 3.6 LC-NE neurons exhibits an essential dependency within the DR-LC-PBC neural circuit, synergistically interacting to regulate S-IRA

Above results have shown that DR-LC neural circuit could reduce the incidence of S-IRA, while previous research results have shown that activating the DR-PBC neural circuit has effectively inhibited the S-IRA, but the significance of the LC as a relay station is unclear. To further verify whether the DR and LC exhibit an interdependent relationship in jointly regulating SUDEP, we injected virus and optical fibers into the DR and DSP-4 was continuously administered intraperitoneally for 1d or 7d during the subsequent week (Fig.9A-C). The results showed that after optogenetic activation of DR5-HT neurons, the incidence of S-IRA with photostimulation was significantly decreased, however, the inhibitory effects could be reversed by DSP-4, which could degrade central LCNE neurons (Fig.9D-E). This suggests that activation of DR5-HT neurons, which depends on the LC-NE neurons, exerts its effect in reducing the incidence of S-IRA. Byword, the 5-HTergic and NEergic systems intrinsically regulate the SUDEP in a synergistic-dependent manner.

**Figure 9.**
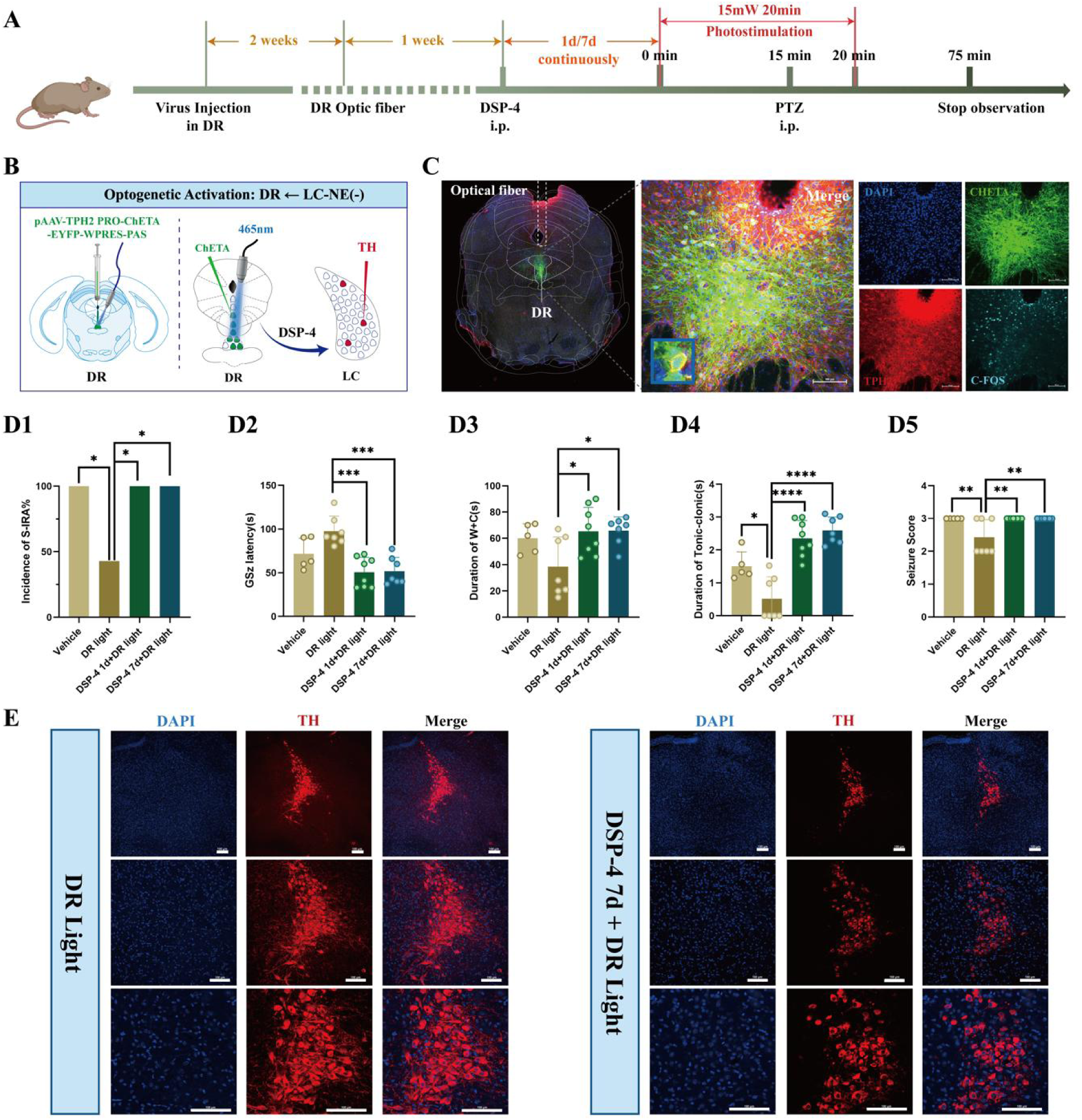
Optogenetic activation of DR 5-HT neurons reduced the incidence of S-IRA evoked by PTZ, and DSP-4 which degraded central LC NE neurons can significantly reverse the effect. **(A)** Experimental protocol for optogenetic activation of DR 5-HT neurons and IP injection of DSP-4 in the PTZ-induced TH-Cre DBA/1 mice. **(B)** Schematic diagram of the positions for the virus (pAAV-TPH2 PRO-ChETA-EYFP-WPRES-PAS) injection and optic fibers implantation in DR. **(C)** Representative images show the optical fiber location and the co-expression of ChETA, C-FOS, TPH2 and DAPI in the DR. **(D1)** The incidence of S-IRA evoked by PTZ, the duration of tonic-clonic seizures and the seizure score in the group with DR light was markedly reduced compared to the group with Vehicle, the group with DSP-4 1d + DR light and the group with DSP-4 7d + DR light. The GSz latency in the group with DR light was significantly higher than the group with DSP-4 1d + DR light and the group with DSP-4 7d + DR light. The duration of wild running and clonic seizures in the group with DR light was markedly lower than the group with DSP-4 1d + DR light and the group with DSP-4 7d + DR light. **(E)** Representative images of staining for TH and DAPI in the LC in the groups with DR light and the group with DSP-4 7d + DR light. ****P < 0.0001; ***P < 0.001; **P < 0.01; *P < 0.05; Data are mean±SEM; DR Light: DR photostimulation; i.p.: intraperitoneal injection

### 3.7 DR-LC-PBC neural circuit collectively regulates SUDEP in a synergistic-dependent manner through the cooperation of central 5-HTergic and NEergic systems by 5-HT2A and NE-α1 receptors

To determine the DR-LC-PBC neural projection and 5-HT2A or NE-α1 receptor responsible for exerting the aforementioned effects, we combined optogenetic approaches with pharmacology, by injecting Prazosin through cannulas within the PBC to block NE-α1 receptors and simultaneously activating the bilateral LC by photostimulation to further observe the effects of both related SUDEP (Fig.10A). AAV-EF1a-DIO-hChR2 (H134R)-eYFP was injected into the bilateral LC and optical fibers were implanted while drug-delivery cannulas were embedded into the bilateral PBC (Fig.10B-C). We found that photostimulation of LCNE neurons showed a significantly lower incidence of S-IRA evoked by PTZ-injection compared with the other groups, which was significantly reversed after the microinjection of NE-α1 receptor antagonist Prazosin into the PBC. Similar results were also found in the duration of wild running, the duration of clonic seizure or tonic-clonic seizures and seizure score (Fig.10D).

**Figure 10.**
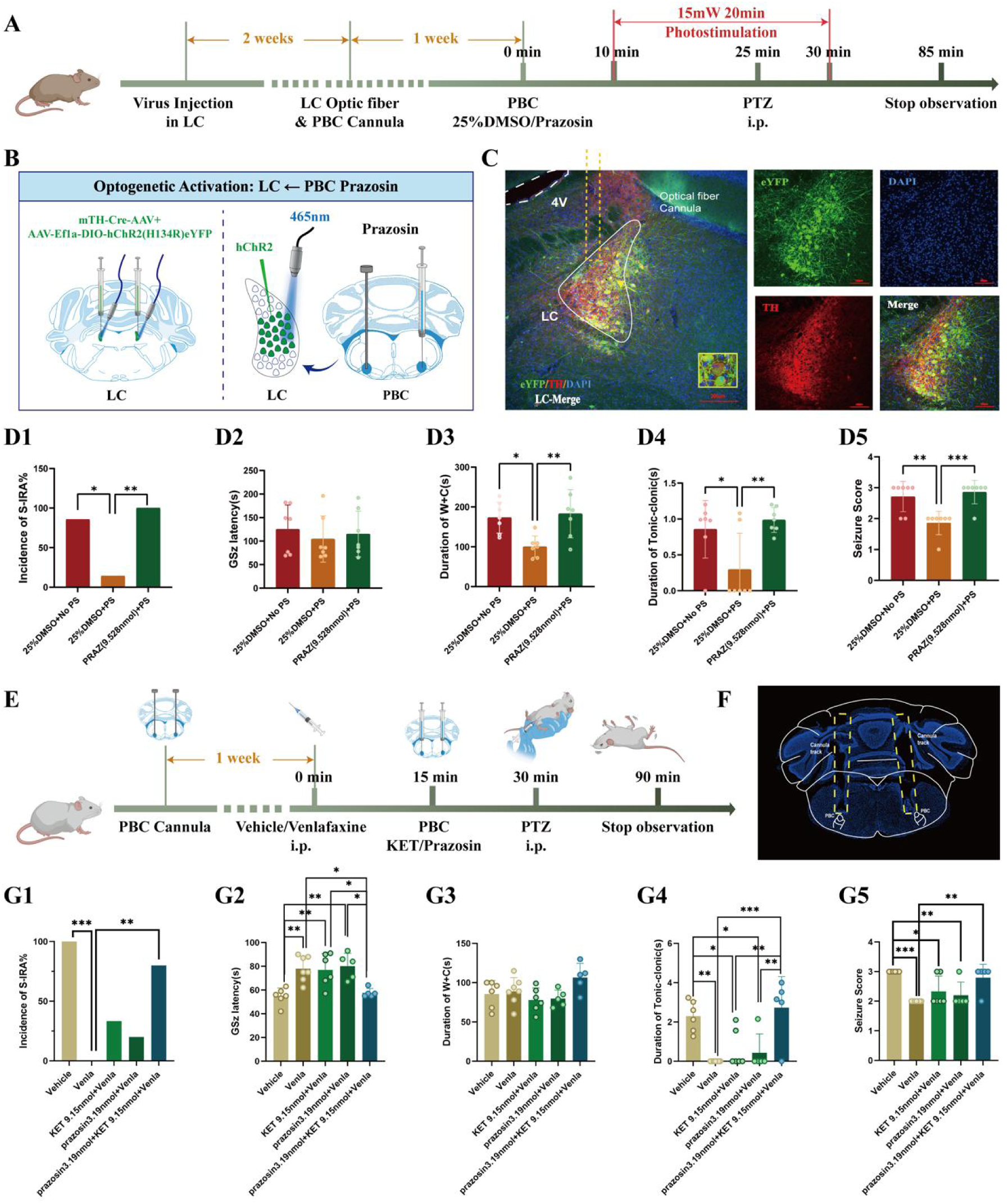
The inhibitory effects of activation of DR-LC-PBC neural circuit can be reversed by 5-HT2A and NE-α1 receptor antagonists microinjected in PBC. **(A)** Experimental protocol for optogenetic activation of LC NE neurons and microinjection of NE-α1 receptor antagonist Prazosin into bilateral PBC in the PTZ-induced TH-Cre DBA/1 mice. **(B)** Schematic diagram of the positions for virus (mTH-Cre-AAV+AAV-EF1a-DIO-hChR2(H134R)-eYFP) injection and optic fibers in the bilateral LC and Prazosin microinjection into the bilateral PBC. **(C)** Representative images show the optical fiber locations and the co-expression of eYFP and TH in the LC. **(D1-5)** The incidence of S-IRA evoked by PTZ, the duration of wild running and clonic seizure or tonic-clonic seizures in the group with 25% DMSO + PS was significantly reduced compared to the group with 25% DMSO + NO PS and the group with PRAZ + PS. The seizure score in the group with 25% DMSO + PS was significantly lower compared to the group with 25% DMSO + NO PS and the group with PRAZ + PS. No obvious differences were observed between treatment groups in the GSz latency (*p* > 0.05). **(E)** Experimental protocol for embedding the cannulas in the bilateral PBC, IP injection of Venlafaxine, and microinjecting 5-HT2A receptor or NE-α1 receptor antagonists into the bilateral PBC in the PTZ-induced DBA/1 mice. **(F)** Representative image shows the tracks of cannulas implanted into the bilateral PBC. **(G1-5)** The incidence of S-IRA evoked by PTZ in the group with Venlafaxine was significantly reduced compared to the group with Vehicle and the group with Prazosin 3.19nmol + KET 9.15nmol + Venlafaxine. The GSz latency in the group with Venlafaxine, the group with KET 9.15nmol + Venlafaxine and the group with Prazosin 3.19nmol + Venlafaxine was markedly higher than the group with Vehicle and the group with Prazosin 3.19nmol + KET 9.15nmol + Venlafaxine. The duration of tonic-clonic seizures in the group with Venlafaxine, the group with KET 9.15nmol + Venlafaxine and the group with Prazosin 3.19nmol + Venlafaxine was markedly lower than the group with Vehicle, and the same significant goes for the group with Venlafaxine, the group with KET 9.15nmol + Venlafaxine and the group with Prazosin 3.19nmol + Venlafaxine. The seizure score in the group with Venlafaxine, the group with KET 9.15nmol + Venlafaxine and the group with Prazosin 3.19nmol + Venlafaxine was markedly lower than the group with Vehicle. The seizure score in the group with Venlafaxine was significantly reduced compared to the group with the group with Prazosin 3.19nmol + KET 9.15nmol + Venlafaxine. No obvious differences were observed between treatment groups in the duration of wild running and clonic seizures (*p* > 0.05). ***P < 0.001; **P < 0.01; *P < 0.05; Data are mean±SEM; PS: photostimulation; NO PS: no photostimulation; PRAZ: Prazosin; i.p.: intraperitoneal injection

To further verify the role of 5-HT2A and NE-α1 receptors involved in regulation of S-IRA, cannulas were embedded in the bilateral PBC and 5-HT2A or NE-α1 receptor antagonists were injected into the PBC after intraperitoneal injection of Venlafaxine (Fig.10E-F). We found that microinjection of a 5-HT2A or NE-α1 receptor antagonists into the PBC can reverse the pro-arousal effect of intraperitoneal injection of Venlafaxine, and the effect of Venlafaxine in inhibiting S-IRA was almost completely reversed while both receptors were simultaneously blocked (Fig.10G), further suggesting that the 5-HT2A and NE-α1 receptors in this circuit participated in regulating SUDEP through the synergistic-dependent cooperation of central 5-HTergic and NEergic systems.

## 4 Discussion

SUDEP is the leading cause of mortality among epilepsy patients and a pressing public health concern. Research on the mechanisms and potential contributing factors of SUDEP has progressed, due to the complexity of SUDEP and the difficulty in obtaining real-time clinical data, the pathophysiological mechanisms remain unclear. Our research group has long been investigating the mechanisms by which the 5-HT and NE neurotransmitter systems affect the occurrence and development of SUDEP. As previous studies have shown, exogenously increasing 5-HT levels in DBA/1 mice and optogenetic activating 5-HT neurons in the DRN can significantly reduce the incidence of S-IRA and SUDEP in mice, and there is a neural circuit connection with the pre-Bötzinger complex (PBC), which regulates respiratory rhythm^11,27,35^. It has also been found that the norepinephrine reuptake inhibitor Atomoxetine can significantly reduce the incidence of S-IRA and SUDEP in DBA/1 mice by increasing NE content in the brain, with the LC likely being an important target of Atomoxetine^17,21^.

The 5-HT and NE systems are important members of the monoaminergic neurotransmitter family, and we deduce that there is an internal regulatory manner within the monoaminergic neurotransmitter system, with some manners between the 5-HT and NE systems that jointly regulates SUDEP. However, the specific regulatory roles, target sites, and whether there is a temporal relationship between the two systems remain unclear. Therefore, we conducted a series of studies using pharmacology, fiber photometry, optogenetics, and chemogenetics to explore the synergistic regulatory manner between the 5-HT and NE systems related to SUDEP.

To further explore the role of the 5-HT and NE systems in the pathogenesis of SUDEP, we respectively constructed acoustic stimulation-induced and PTZ injection-induced SUDEP models. Injection of 5-HTP or Atomoxetine alone to elevate 5-HT and NE levels revealed that a low dose of 100mg/kg 5-HTP or 5mg/kg Atomoxetine did not reduce the incidence of S-IRA and SUDEP. Nevertheless, at effective doses of 125mg/kg 5-HTP or 20mg/kg Atomoxetine, although the incidence of SUDEP in mice was reduced, adverse effects such as diarrhea and poor motor performance were observed, which may be related to the toxic reactions caused by excessive drug doses^36^. To avoid the adverse reactions caused by high drug doses, DBA/1 mice were given IP injections of the aforementioned ineffective low doses of both 5-HTP and Atomoxetine. It was found that compared with the IP injection of effective high doses of 5-HTP or Atomoxetine alone, this significantly and completely blocked the incidence of S-IRA induced by acoustic stimulation and PTZ injection. Moreover, DBA/1 mice at this dose showed no adverse drug reactions and significantly prolonged GSz latency, reduced duration of W+C, duration of tonic-clonic seizures, and seizure score. This indicates that increasing the levels of 5-HT and NE can reduce the incidence of S-IRA and SUDEP, and that the two have some synergistic effect.

Similar to our findings, previous studies have shown that 5-HTP, at specific dosages, can effectively reduce incidence of seizure, decrease frequency, and prolong latency. Moreover, 5-HT levels are intimately linked to the seizure threshold; lowering 5-HT levels diminishes the threshold, thereby exacerbating generalized seizure activity ^37–39^. Concurrently, endogenous NE exhibits anticonvulsant properties in epilepsy. Furthermore, decreased NE levels are correlated with heightened susceptibility to neuronal damage in regions marginally free of seizures and/or prone to audiogenic seizures^40^. The neuroprotective effects of NE are ascribed to its role in counteracting epileptogenic circuit formation and modulating epilepsy-associated neuronal alterations, which collectively reduce central nervous system susceptibility to epileptiform activities^39^. Consequently, the regulation of the 5-HT and NE systems does not merely sum their effects but rather produces a synergistic outcome greater than the sum of their parts.

To further investigate the underlying internal mechanisms, we turned our attention to the representative nuclei—the DRN, the primary source of forebrain 5-HT innervation, and the LC, the major nucleus for NE synthesis and release in the brain. We conducted a series of studies on the DRN-LC neural circuit and its coordinated regulation of SUDEP. We first selected Venlafaxine for pharmacological experiments, which can simultaneously block the reuptake of 5-HT and NE neurotransmitters, thereby increasing the levels of 5-HT and NE. In SUDEP models induced by acoustic stimulation and PTZ injection, we found that IP injection of Venlafaxine reduced the incidence of S-IRA and decreased mortality. In the PTZ-induced SUDEP model, IP injection of Venlafaxine also reduced the duration of tonic-clonic seizures and seizure scores, with no observed behavioral abnormalities in mice during the experiments. Additionally, injection of either PCPA or DSP-4 alone could reverse the effect of Venlafaxine in reducing S-IRA. Combined administration of PCPA and DSP-4 completely blocked the effect of Venlafaxine, shortening GSz latency and increasing duration of W+C and duration of tonic-clonic seizures. Therefore, Venlafaxine can regulate SUDEP through both the 5-HT and NE systems. After Venlafaxine injection, administration of the 5-HT2A receptor antagonist KET and the NE-α1 receptor antagonist Prazosin also reversed the effect of Venlafaxine in reducing S-IRA. Venlafaxine, as a SNRI, epitomizes the synergistic regulation of the 5-HT and NE systems, indicates that the 5-HT system associated with the DRN, and the NE system linked to the LC can jointly modulate the SUDEP, with this effect mediated through 5-HT2A and NE-α1 receptors.

Notably, after IP injection of Venlafaxine, immunohistochemistry revealed a significant increase in c-fos activity of DRN^5-HT^ neurons and LC^NE^ neurons. With increasing doses of Venlafaxine, the activity of these neurons was suppressed, a trend consistent with behavioral changes following IP Venlafaxine injection. This indicates that during the regulation of SUDEP, both DRN^5-HT^ neurons and LC^NE^ neurons are involved, showing an initial activation followed by suppression. Subsequent ICV injection of Venlafaxine and calcium signal fiber recordings of DRN^5-HT^ neurons and LC^NE^ neurons revealed that varying doses of Venlafaxine led to increased or decreased incidence of S-IRA, corresponding to the activation or inhibition of these neurons. This strongly suggests a close synergistic relationship between DRN^5-HT^ and LC^NE^ neurons in the regulation of SUDEP.

The descending projections from the DRN to the LC account for at least 50% of the 5-HT innervation of this nucleus. The LC also receives afferent neurons from the median raphe nucleus, and the 5-HT neurons in the DRN are densely innervated by NEergic neurons from the LC. Inhibition of DRN activity can significantly reduce the release of NE in the LC. To investigate the DRN^5-HT^ neurons and LC^NE^ neurons, we employed optogenetic and chemogenetic techniques. It was found that optogenetic activation of DRN^5-HT^ neurons significantly reduced the incidence of S-IRA and activated LC^NE^ neurons. Similarly, optogenetic activation of LC^NE^ neurons significantly reduced the incidence of S-IRA but did not significantly activate DRN^5-HT^ neurons, a finding also confirmed by chemogenetic activation of LC^NE^ neurons. Therefore, we hypothesize that DRN5-HT neurons may act upstream to regulate downstream LCNE neurons, which is a synergistic manner coordinating 5-HTergic and NEergic system involvement in SUDEP.

As our previous research on the mechanisms of DRN-associated 5-HTergic and LC-associated NEergic systems in regulating SUDEP has shown, the DRN and LC are pivotal relay points for the synthesis, release, and transmission of neurotransmitter signals in the monoaminergic neurotransmitter system, but they are not the final determinants in regulating SUDEP. Our extensive previous research indicates that S-IRA can induce SUDEP in DBA/1 mouse models through acoustic stimulation or PTZ injection^21,35,41^, aligning with the mainstream view that S-IRA is a primary factor in SUDEP. One critical finding of our research is the involvement of the 5-HTergic neural circuit between the DR and PBC in S-IRA and SUDEP. Previous studies have also shown that there is a neural circuit between the LC and PBC that controls respiration^42^

Given the pivotal role of the PBC in regulating respiratory rhythm, our focus shifted to the PBC, identifying it as the optimal candidate for the final relay point where DR-related serotonergic and LC-related noradrenergic systems synergistically regulate SUDEP. To further investigate the bridges among the DR, LC, and PBC and their roles in regulating SUDEP, we utilized anterograde tracing virus HSV and retrograde tracer CTB-555 for dual confirmation, establishing the neural anatomical circuit among the DR, LC, and PBC. Injection of anterograde tracing virus HSV into DRN^5-HT^ neurons revealed that the majority of bilateral LC^NE^ neurons and bilateral PBC neurons were labeled. Similarly, injection of retrograde tracer CTB-555 into bilateral PBC showed that most bilateral LC^NE^ neurons and DRN-5-HT neurons were labeled. Based on previous and current research findings, DRN^5-HT^ neurons, LC^NE^ neurons, and PBC neurons form a hierarchical relationship, with DRN^5-HT^ neurons potentially reducing the incidence of S-IRA by simultaneously activating LC^NE^ and PBC neurons.

However, previous research results have shown that activating the DRN-PBC neural circuit has effectively reduced the S-IRA and SUDEP, prompting us to consider the significance of the LC as a relay station. Optogenetic activation of the DRN-LC neural circuit and chemogenetic activation of LC^NE^ neurons both significantly reduce the incidence of S-IRA, markedly prolong the latency, decrease the duration of tonic-clonic seizures, and lower seizure scores, indicating that the LC is a sufficient condition for regulating the SUDEP. To verify that the LC is a necessary condition for this circuit to regulate SUDEP, we pretreated with a highly selective neurotoxin DSP-4 to degrade central LC^NE^ neurons before optogenetic activation of DRN^5-HT^ neurons. Results showed that DSP-4 significantly reversed the effect of optogenetic activation of the DRN in reducing S-IRA, shortened the latency, and increased the duration of W+C, and the duration of tonic-clonic seizures, indicating that the effect of DRN^5-HT^ neuron activation in reducing S-IRA incidence depends on the LC^NE^ neurons. More importantly, this suggests that the 5-HTergic and NEergic systems intrinsically regulate the SUDEP in a synergistic and interdependent manner.

To further verify the presence of a projection circuit between LC^NE^ neurons and the PBC that regulates the S-IRA and SUDEP, we microinjected an NE-α1 receptor antagonist into the PBC before optogenetic activation of LC^NE^ neurons. The results specifically showed that antagonizing the PBC NE-α1 receptor reversed the effects of optogenetic activation of LC^NE^ neurons, significantly increasing the incidence of S-IRA, duration of W+C, and the duration of tonic-clonic seizures. These findings indicate that the LC is a crucial component in the DRN-LC neural circuit that mediates the influence of the PBC on respiratory rhythm and regulation of S-IRA, making it both a sufficient and necessary condition for this neural circuit to regulate SUDEP. Based on previous findings, we concurrently enhanced 5-HT and NE neurotransmission while intra-PBC injection of KET and Prazosin, revealed that 5-HT2A and NE-α1 receptors are likely the targets through which the 5-HT and NE systems exert their synergistic-dependent effects.

In summary, conclusive evidence is provided in our study that the 5-HT neurotransmitter systems synergistically regulate SUDEP with NE neurotransmitter systems and strongly necessitate the involvement of the NE neurotransmitter system. The results of this study, derived from our long-term SUDEP-related research series, though we do not rule out the possibility that the 5-HT and NE systems may regulate SUDEP through other manner. The 5-HT and NE systems regulated SUDEP through an intrinsic synergistic-dependent manner is the first proposal within the monoaminergic systems in the field of SUDEP pathogenesis research. This stands in contrast to previous studies focusing on single-system effect on SUDEP, providing novel directions for accurately interpreting the mechanisms of SUDEP and offering innovative prevention strategies and intervention targets.

## Authors’ contributions

H.H.Z. contributed to the conceptualization and design of the study. Q.X, X.X.X, Y.X.W. and Y.Y. performed the experiments, analyzed most of the experiments and wrote the draft of the manuscript. Z.W.Z. and X.Y.D. helped with the analysis of the experiments and participated in the modification. All authors contributed to the final version of the manuscript.

## Acknowledgments

We thank YuDong Zhou and Yi Shen for their help in experimental design.

## Data Availability

The data supporting the findings of this study are available within the article. Data will be made available upon reasonable request.

## Declarations

### Ethics Approval

All procedures were in accordance with the National Institutes of Health Guidelines for the Care and Use of Laboratory Animals and were approved by the Animal Advisory Committee of Zhejiang University.

### Consent to Participate

Not applicable.

### Consent for Publication

Not applicable.

## Supplementary materials

**Table S1.**
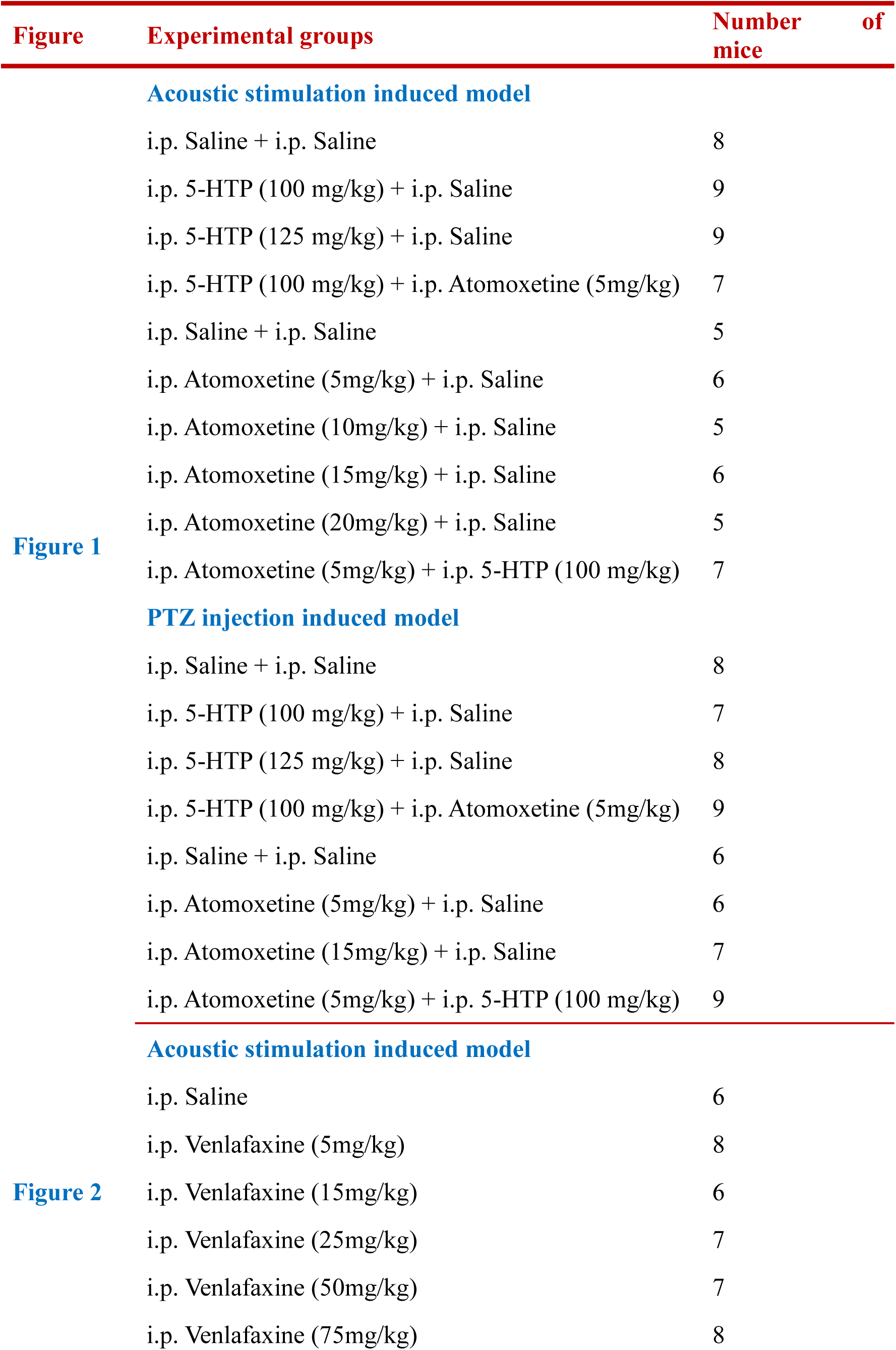
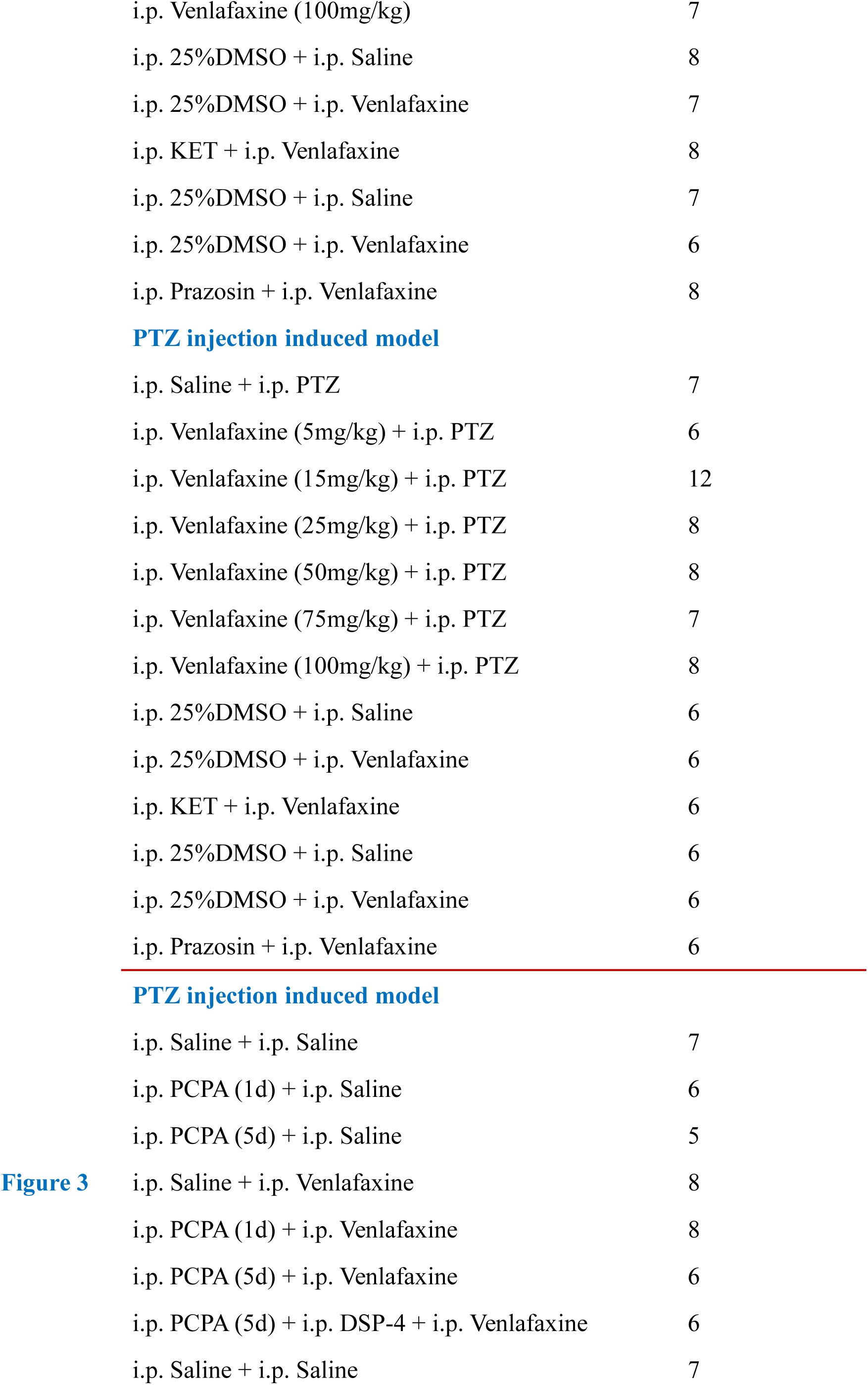
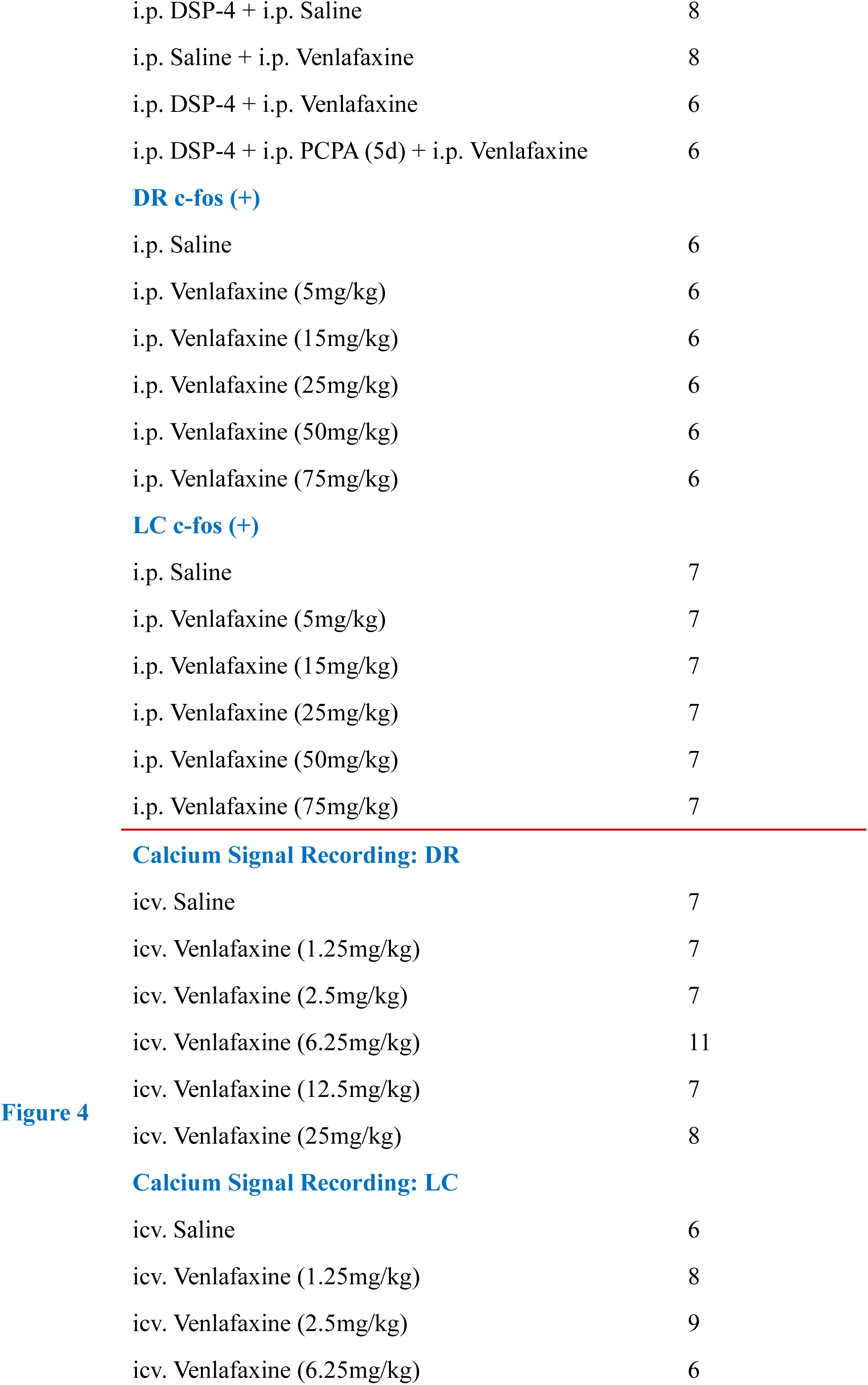
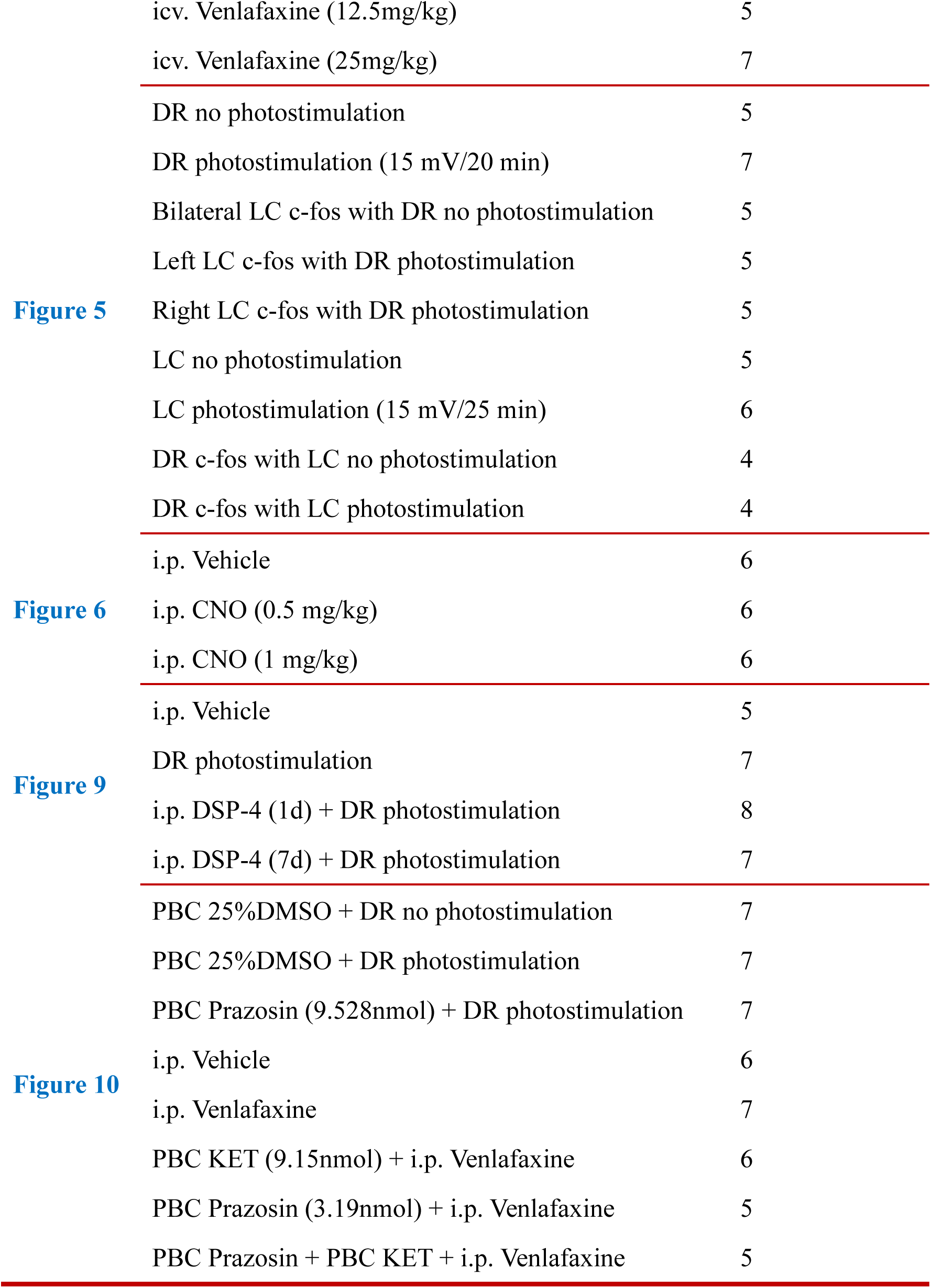
Summary of experimental groups of DBA/1 mice.

**Table S2.**
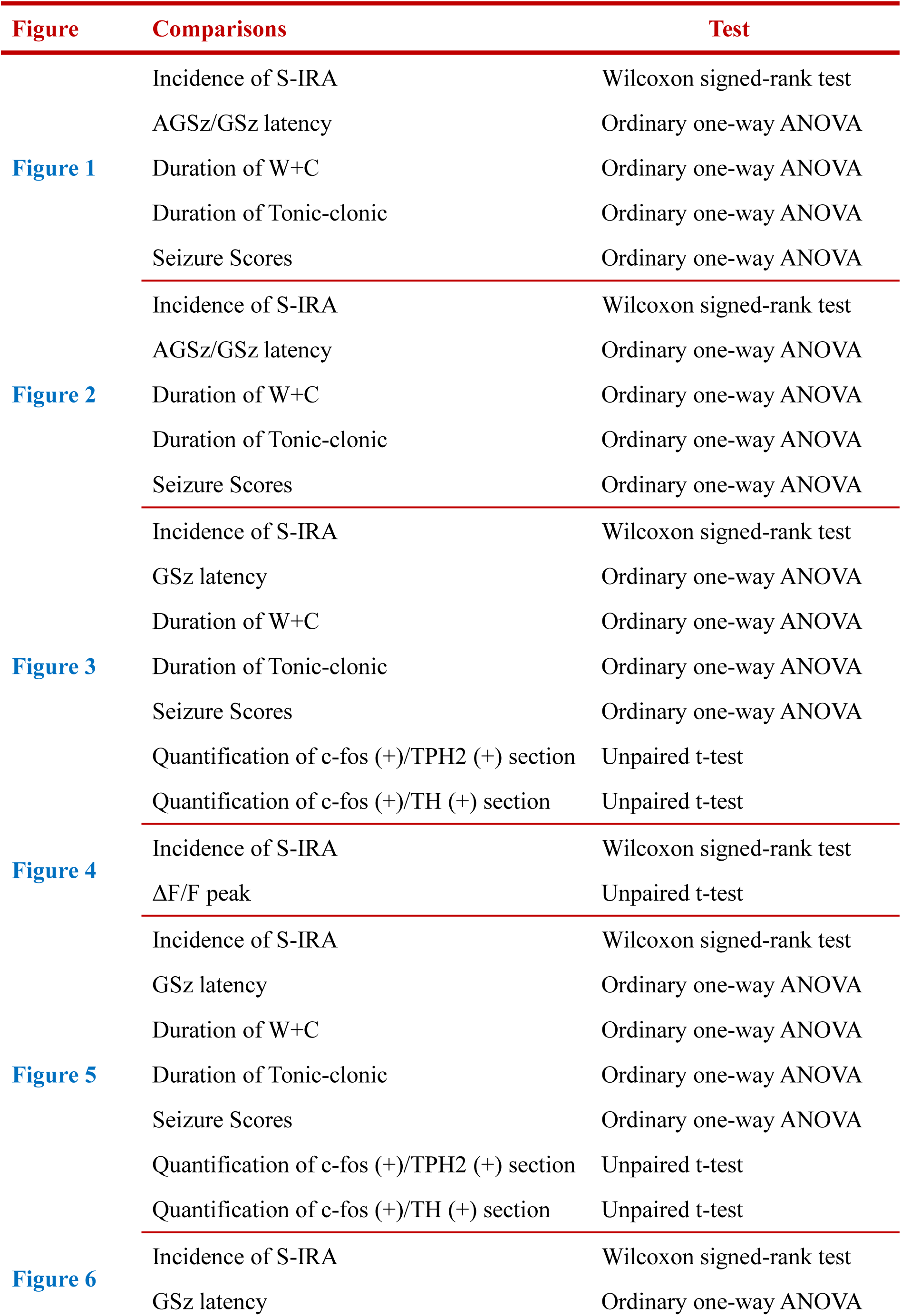
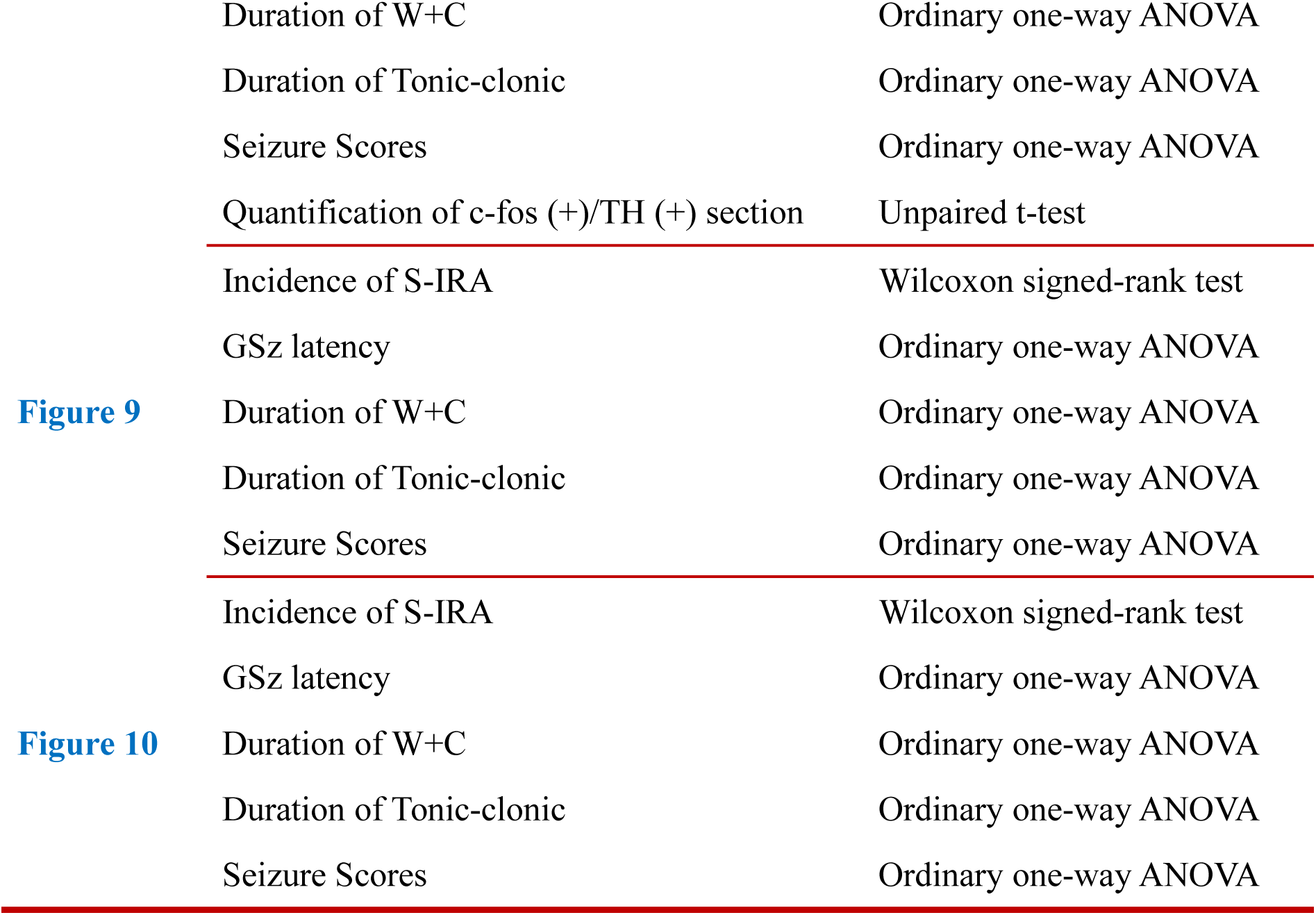
Statistical analysis in each group.

**Table S3.** Reagent or resource.

| Reagent or resource | Source | Identifier |
| --- | --- | --- |
| **Antibodies** |  |  |
| Mouse anti-TH | Merck-Millipore | MAB318 |
| Mouse anti-TPH2 | Sigma-Aldrich | T0678 |
| Rabbit anti-c-fos | Cell Signaling Technology | 2250S |
| Donkey anti-mouse Alexa 546 | Thermo Fisher Scientific | A10036 |
| Donkey anti-mouse Alexa 488 | Thermo Fisher Scientific | A21202 |
| Goat anti-rabbit Cy5 | Thermo Fisher Scientific | A10523 |
| **Bacterial and virus strains** |  |  |
| AAV2/9-mCaMKIIa-GCaMP6f-WPRE-pA | Brain VTA Technology Co., Ltd. | N/A |
| rAAV-DBH-GCaMP6m-WPRE-hGH pA | Brain VTA Technology Co., Ltd. | N/A |
| pAAV-TPH2 PRO-ChETA-EYFP-WPRES-PAS | Shengbo, Co., Ltd. | N/A |
| pAAV-CAG-DIO-CHETA-EGFP | Shengbo, Co., Ltd. | N/A |
| mTH-Cre-AAV+<br>AAV-EF1a-DIO-hM3Dq-mCherry | Brain VTA Technology Co., Ltd. | N/A |
| HSV-1 H129 | Brain VTA Technology Co., Ltd. | N/A |
| mTH-Cre-AAV+<br>AAV-EF1a-DIO-hChR2 (H134R)-eYFP | Brain VTA Technology Co., Ltd. | N/A |
| **Chemicals, peptides, and recombinant proteins** |  |  |
| PTZ | Sigma-Aldrich | Cat #P6500 |
| 5-HTP | Sigma-Aldrich | Cat #107751 |
| Atomoxetine | Sigma-Aldrich | Ca #Y0001586 |
| Venlafaxine | Sigma-Aldrich | PHR1736 |
| DSP-4 | Sigma-Aldrich | C8417 |
| PCPA | Sigma-Aldrich | C3635 |
| KET | Sigma-Aldrich | Cat #8006 |
| Prazosin | Sigma-Aldrich | Cat # P7791 |
| DMSO | Sigma-Aldrich | SHBK2703 |
| Clozapine N-oxide (CNO) | Brain VTA Technology Co., Ltd. | N/A |
| CTB-555 | Brain VTA Technology Co., Ltd. | N/A |
| **Experimental models: Organisms/strains** |  |  |
| TH-Cre C57BL/6J mice | Jackson Lab | B6.Cg-7630403G23RikTg (TH-Cre) 1Tmd/J |
| **Software and algorithms** |  |  |
| Fiber photometry system | Inper | C11946 |
| ImageJ | NIH, Bethesda, MD, USA | https://imagej.nih.gov/ij/ |
| GraphPad Prism TM8.0 |  |  |
| SPSS version 23.0 |  |  |

